# Visual surfaceomics reveals the substrate-scavenging cell surface of Asgard archaea

**DOI:** 10.64898/2026.11.05.763656

**Authors:** Jessie James Limlingan Malit, Davide Amendola, Jingwei Xu, Laura Dufresne, Anastasiia Kokhanovska, Antonina Gonet, Charlotte P. Wiegand, Julien Clavien, Florian Wollweber, Martin Pilhofer

## Abstract

The emergence of eukaryotes from Asgard archaeal ancestors^1–5^ likely required a specialized cell envelope that enabled sustained physical and metabolic interactions with proto-mitochondrial symbionts^6–10^. While the protrusion-rich cell architecture of Asgard archaea may enable such interactions, the molecular composition of their cell surface and its role in establishing cell-cell contacts remains unknown.

Here, we developed a “visual surfaceomics” approach, integrating subtomogram averaging, predictive structural modeling, transcriptomics, proteomics, and expansion microscopy to identify surface proteins *in situ*. We show that Asgard cell bodies and protrusions lack an archaea-typical paracrystalline S-layer and are instead decorated by two main classes of surface proteins: subtilisin-like serine proteases (SubSs) and oligopeptide-binding proteins (OppAs). SubSs were detected as membrane-anchored dimers and were sparsely distributed across the surface. OppAs showed near-total membrane coverage but lacked any rigid, higher order assembly. Thermal shift assays and single particle cryo-EM showed that OppA binds oligopeptides *in vitro*. Comparative genomics, transcriptomics, and proteomics suggest that this peptidophilic surface architecture is conserved across Asgard archaeal phyla. Finally, we discovered that the cell surface was remodeled at cell-cell contact sites.

Altogether, our integrative approach revealed a metabolically specialized and structurally flexible cell surface that couples extracellular protein degradation with peptide capture and uptake, while potentially contributing to establishing and maintaining the symbiotic interactions of Asgard archaea.

## Introduction

Eukaryotes are assumed to have emerged from an endosymbiosis between an archaeon and the alpha-proteobacterial mitochondrial ancestor^6,7^. Although several models of this process have been postulated, they generally agree that the proto-mitochondrion initially formed a close association with the archaeon for metabolic exchange^1,7–9,11^. Mechanistically, this may have required the remodeling or weakening of the rigid archaeal cell surface, or potentially its loss, to facilitate closer interactions with the symbiont^9^. These changes could have also enabled the formation of membranous protrusions, thereby increasing the surface area for enhanced symbiotic interactions^1,9,10^ and potentially leading to a stepwise protrusion-mediated engulfment of the proto-mitochondrion, forming the first eukaryotic cell.

The archaeal host is likely related to Promethearchaeati (here referred to as Asgard archaea), based on phylogenetic analyses and the extensive repertoire of eukaryotic signature proteins (ESPs) in their genomes^2–5^. Several members of the Asgard archaea have recently been cultivated, all of which require the presence of one or more syntrophic partners, usually a methanogen^1,12^ or sulfate-reducing bacteria^1,13,14^ for growth and survival. Imaging of these cells consistently showed cell bodies connected to protrusion networks and membrane-bound vesicles^1,12–15^. In some cases, potential cell-cell interactions with the other species in the enrichments were also observed^13,14^. This complex morphology could potentially be formed and stabilized by their eukaryote-like cytoskeleton composed of abundant actin filaments and, more rarely, microtubules^13,16^. The Asgard archaeal cell envelope composition and architecture however, remains unknown.

Most archaea possess a paracrystalline, proteinaceous S-layer that envelops their cell membrane, imparting rigid support and osmoprotection, and enabling the transport of metabolites and interactions with other cells^17,18^. The lokiarcheal strain *Prometheoarchaeum syntrophicum* and the hodarchaeal strains *Ca.* Margulisarchaeum peptidophila and *Ca.* Flexiarchaeum multiprotrusionis have been suggested to possess a canonical S-layer-like structure composed of symmetrically arranged homopolymers^1,12^. In contrast, the surfaces of the Lokiarchaea *Ca.* L. ossiferum and *Nererarchaeum marumarumayae*, and the Heimdallarchaeon *Ca.* Yibarchaeum umbracryptum were described to have a heterogeneous and irregular proteinaceous layer that contrasts with the typical S-layer of archaea^13–15^. Small and unordered densities were found covering most of their cell surfaces, but more elaborate and complex membrane-bound structures were also present. These diverse proteins were observed to e.g. connect separate protrusions or localize to sites of high membrane curvature^13^. Previous bioinformatic and glycoproteomics studies have proposed the identity of some Asgard surface proteins, but none of these predictions have been experimentally validated^1,12,14^. Uncovering the identity and function of this assemblage of surface proteins found in Asgard archaea could illuminate the interactions with their syntrophic partners and the environment as a proxy for the cell-cell interactions at the onset of eukaryogenesis.

## Results

### Lokiarchaeal cell envelopes are covered with irregularly arranged and elaborate cell surface densities

To systematically characterize the cell surface of Lokiarchaea, we leveraged our cryo-tomography dataset of *Ca.* L. ossiferum (57 tomograms^16^) plus one additional tomogram at higher magnification (Fig. 1a), and acquired a new dataset for *Prometheoarcheum syntrophicum* (99 tomograms) and its syntrophic partner *Methanogenium sp.* (8 tomograms). Consistent with previous observations, both lokiarchaeal species exhibit pleomorphic cellular morphologies (Fig. 1a-d, Extended Data Fig. 1a,c) with cell bodies connected to numerous protrusions. Notably, the cell membranes of both species were decorated with irregularly arranged and variably sized densities found in both cell bodies and protrusions. This markedly differs from an archaeal S-layer-like organization, as observed in *Methanogenium sp.* (Fig. 1e). Two distinct classes of densities could be discerned: “small surface densities (SSDs)” and “large surface densities (LSDs).” The SSDs protrude uniformly to around ∼5.5 nm from the cell membrane with a length of ∼7 nm along their longest axis (Extended Data Fig. 1b). The LSDs extend to ∼25 nm from the membrane with a width of up to ∼16 nm (Extended Data Fig. 1b). These large densities consist of two lobes emanating from a stem-like structure (insets in Fig. 1a,c).

**Figure 1.**
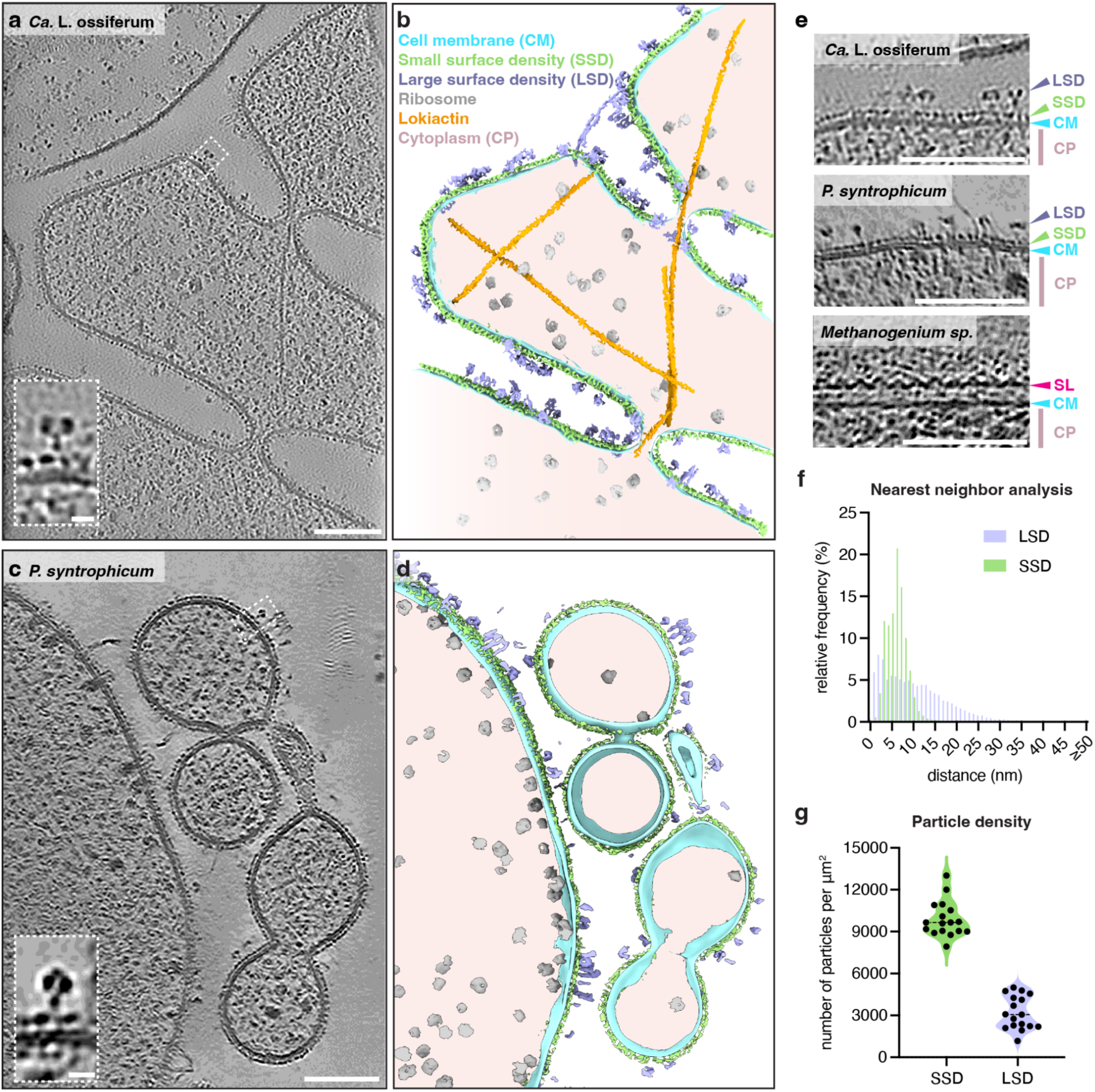
Lokiarchaeal cell envelopes are covered with irregularly arranged and elaborate cell surface densities. **a**-**d**, Slices through cryo-tomograms (**a,c**) and corresponding segmentations (**b,d**) of *Ca.* L. ossiferum (**a,b**) and *P. syntrophicum* (**c,d**). Insets are enlarged views of boxed areas, showing the large (LSD) and small (SSD) surface densities found on the cell surface of Lokiarchaea. Scale bars: 100 nm and 10 nm for enlarged views. **e**, Slices of cryo-tomograms showing a comparison of the cell envelopes of both Lokiarchaea and a methanogen, which shows an archaea-typical S-layer (SL). Scale bars: 100 nm. **f,g,** Distribution of the two major cell surface densities (Extended Data Fig. 1b) quantified across 17 cryo-tomograms of *Ca.* L. ossiferum. (**f**) shows the calculated geodesic nearest neighbor distance between individual densities and (**g**) shows number of particles per µm^2^ cell membrane area for both classes of surface densities. While SSD shows near-total membrane coverage, LSD shows a sparser and more variable distribution. See also Extended Data Fig. 1a for an example of higher LSD density.

To characterize the distribution and localization of these two classes of surface densities, we employed deep learning-based tools to segment the tomograms^19^ and to extract the coordinates^19,20^ for a subset of *Ca.* L. ossiferum tomograms. We then analyzed their distribution by calculating geodesic nearest neighbour distances of particle centroids projected onto the nearest membrane mesh. The nearest neighbor distances between SSDs averaged 6.05 nm (± 2.62 nm), being consistent with a near total membrane coverage (Fig. 1f, Extended Data Fig. 1e). Owing to the thinner cell bodies and protrusions of some *P. syntrophicum* cells, some tomograms provided particularly insightful top views of the cell surface (Extended Data Fig. 1e), confirming dense but spatially disordered orientations of SSDs across the membrane.

In contrast, the LSDs were on average two to threefold less abundant than the SSDs, resulting in more variable inter-particle distances (*x̅* = 10.45 nm ± 8.04 nm, Fig. 1f,g). The data indicate heterogeneity regarding the distribution of LSDs, ranging from closely packed to being separated by more than 20 nm (Fig. 1f,g, examples with different LSD density are shown in Fig. 1a and Extended Data Fig. 1a). Since no intracellular features could be correlated to the presence of LSDs, and no obvious differences could be observed when comparing cell bodies vs. protrusions, the observed heterogeneity may be based on cell-to-cell variability.

Altogether, the *P. syntrophicum* and *Ca.* L. ossiferum cell surfaces are distinct from the cell surface of most S-layer containing archaea, featuring an irregular assemblage of two major classes of surface densities. This is supported by previous bioinformatic analyses, showing that Asgard archaeal genomes lack homologs of canonical S-layer proteins that could produce a lattice^21^.

### Visual surfaceomics identifies a subtilisin-like serine protease as an abundant lokiarchaeal surface protein

Next, we sought to identify both types of surface densities. Since known lokiarchaeal cultures grow only slowly, to low cell densities, and maintained in non-axenic cultures, conventional approaches, including surface shaving mass spectrometry, surface protein enrichment through biotinylation, and surface glycoprotein purification^22–24^ were not applicable. Subtomogram averaging, an imaging-based approach, has been used previously to characterize surface proteins^25–29^. The thickness of the cells, the lack of lattice organization and the very small size of the observed surface densities, however, prevent achieving high resolution that would allow for *de novo* protein identification. Moreover, the absence of genetic tools prevents validation through conventional deletion or tagging approaches.

We therefore developed an integrative “visual surfaceomics” pipeline to identify and validate the identities of both types of membrane-anchored densities (Fig. 2a-d). Using this pipeline, we set out to firstly identify the protein(s) constituting the LSDs of *Ca.* L. ossiferum. From the whole cell proteomics data, we extracted 152 candidate proteins with predicted N-terminal signal peptides (SP) and/or transmembrane domains (TM), indicative of potential membrane anchoring or secretion. We generated AlphaFold-predicted^30^ structures for each candidate and, in parallel, calculated a representative density map of the LSDs by subtomogram averaging (Extended Data Fig. 2). This yielded a ∼15 Å-resolution reconstruction that reveals a potentially dimeric density consisting of two individual “stalks” extending from the membrane and two globular “head” domains (Fig. 2c). We then systematically fitted each AlphaFold-predicted structure into the putative asymmetric unit of the experimental density map. To this end, we adapted the first step of the integrative structural modeling tool Assembline^31^ to generate fit libraries between coarse-grained, rigid-body representations of each of the AlphaFold structures and the density map. The cross-correlation metric “overlap” was used to quantify the agreement between each predicted structure and the experimental map following random initial placements and global searches^31,32^. Higher overlap scores indicate better agreement between the predicted structure and experimental density. Among the 152 candidates, one protein showed the highest overlap score while also exhibiting high proteomic abundance, strongly supporting its assignment to each of the putative monomers of the LSDs (Fig. 2d), which we further confirmed via visual inspection of the fit (Fig. 2e). We hereafter refer to this protein as LoSubS1 (UYP45189.1). Two copies of LoSubS1 account for the majority of the determined map (Fig. 2e). An additional unresolved density at the base of the reconstruction may correspond to the SSDs or another protein involved in membrane anchoring.

**Figure 2.**
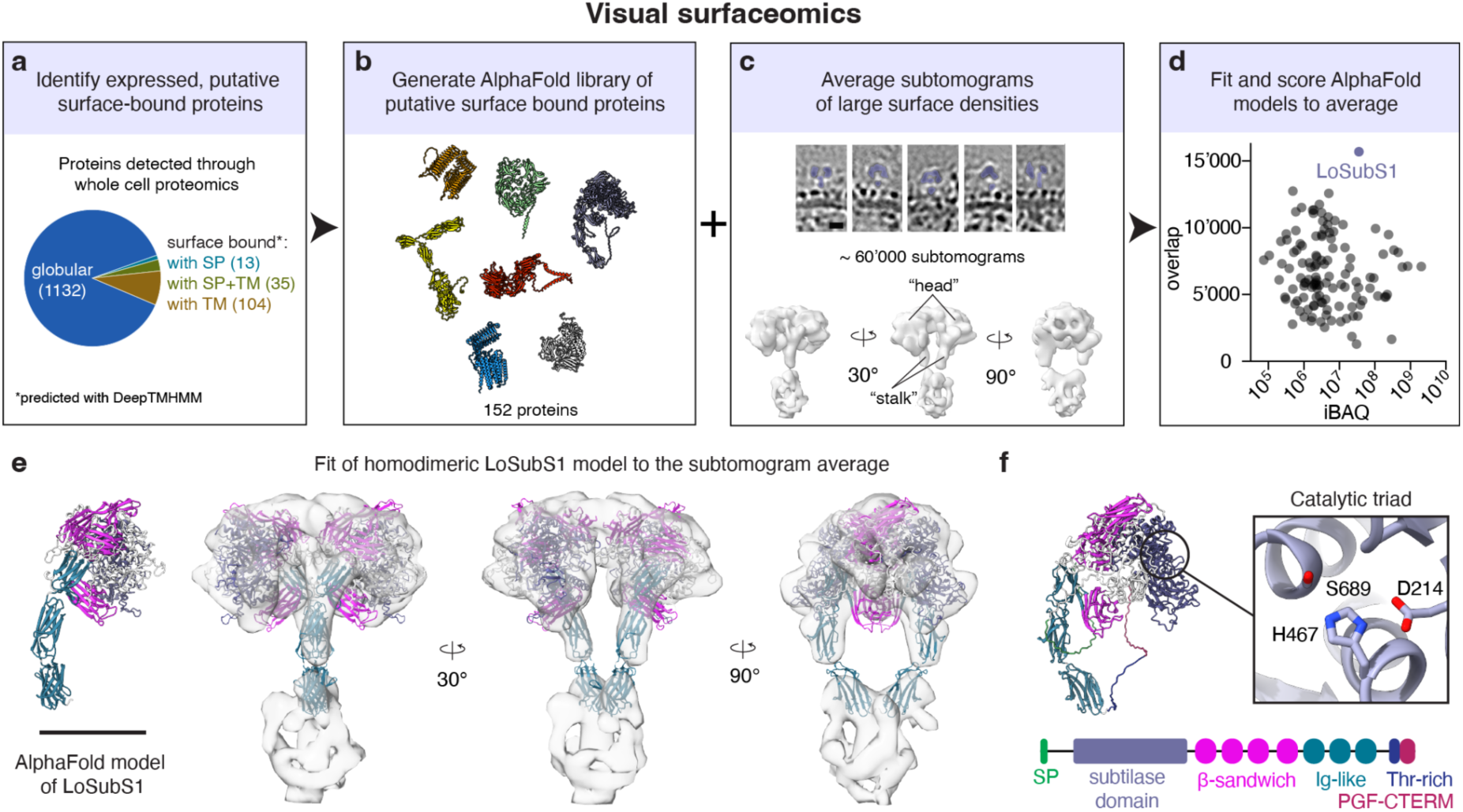
Visual surfaceomics identifies a subtilisin-like serine protease as a major lokiarchaeal surface protein. **a-d,** Visual surfaceomics workflow for the identification of surface densities in cryo-tomograms of Lokiarchaea. First, whole cell proteomics data were filtered for predicted surface-bound or secreted proteins based on the presence of a signal peptide (SP) and/or transmembrane domain/s (TM) (**a**). Then, an AlphaFold library of structures was generated from the filtered putative surface-bound proteins (**b**). A reconstruction of the surface density was calculated through subtomogram averaging using the acquired cryo-ET data (**c**). Shown are representative subvolumes (top) and the subtomogram average (14.8 Å resolution) of the LSDs from *Ca.* L. ossiferum. Lastly, predicted structures of candidate surface-bound proteins were systematically fitted into the putative monomer of the obtained reconstruction (**d**). The overlap score of the reconstruction with each of the predicted structures was plotted against the protein abundances (iBAQ values derived from whole cell proteomics data) of the candidates. LoSubS1 has the highest overlap score as compared to all the other candidate proteins. **e,** Docking of the AlphaFold-predicted structure of LoSubS1 (shown on left, N-terminal SP and C-terminal Thr-rich and PGF-CTERM domains removed) into the reconstructed map (transparent). The map is consistent with the accommodation of a LoSubS1 homodimer. Scale bar: 10 nm. **f,** Predicted structure and domain organization of LoSubS1 based on both sequence and structure homology searches, showing the subtilase domain and structural domains composed of β-sandwiches and Ig-like folds. The protein also contains a signal peptide (SP) and a PGF-CTERM domain, typical features of archaeal surface-anchored proteins. The AlphaFold-predicted structure also shows the conserved catalytic domain (magnified view) of the subtilisin-like serine protease domain.

Functional annotation of LoSubS1 using sequence and structure homology searches identifies a S8-family serine peptidase/subtilase domain (CDD cd07497, E-value = 9.87×10^−41^; PFAM PF00082, E-value = 1.7×10^−22^) (Fig. 2f, Extended Data Fig. 3a). Subtilases are widely distributed across the tree of life^33^ and in archaea, they are anchored to the cell membrane or secreted to the extracellular environment to digest proteinaceous substrates into smaller oligopeptides that can be subsequently used for metabolism^34^. A subtilase domain with an exodigestive role is also present in the S-layer protein of the archaeon *Staphylothermus thermus*^35^. Consistent with a proteolytic function, the AlphaFold model of LoSubS1 contains the characteristic His-Ser-Asp catalytic triad of serine proteases^33^ (Fig. 2f). The predicted structure additionally contains several repeats of β-sandwiches and Ig-like folds, structural motifs commonly found in archaeal S-layer proteins^17,18,25,36^. An N-terminal signal peptide and a C-terminal PGF-CTERM domain (Extended Data Fig. 3b,c) provide a putative mechanism for translocation across the membrane and lipid anchoring of LoSubS1 at the cell surface^37^. A threonine-rich region next to the C-terminal sorting signal of LoSubS1 may provide potential sites for O-glycosylation and could act as a flexible spacer that positions LoSubS1 away from the membrane and the underlying SSDs, as observed in several archaeal S-layer proteins^25^. A homolog of LoSubS1 is also found in the *P. syntrophicum* genome, hereafter referred to as PsSubS1 (QEE15903.1, Extended Data Fig. 3d). Similar to LoSubS1, PsSubS1 contains a subtilase domain alongside several β-sandwiches and Ig-like folds (Extended Data Fig. 3d), and an N-terminal signal peptide and a C-terminal transmembrane domain. Unlike LoSubS1, PsSubS1 lacks a PGF motif and thus is likely anchored to the cell via its C-terminal transmembrane helix.

### Oligopeptide-binding proteins are the major lokiarchaeal cell surface proteins

Since our visual surfaceomics workflow enabled us to identify a candidate protein for one of the major surface densities, we next attempted to uncover the identity of the SSDs. Since these particles are substantially smaller than the LSDs and far below the size of particles typically analyzed by in situ cryo-ET, we applied a more stringent approach during subtomogram averaging (Extended Data Fig. 4a), including manual removal of misannotated particles and assignment of initial particle orientations based on local membrane geometry. This approach yielded an approximately 18 Å-resolution reconstruction (Fig. 3a, Extended Data Fig. 4b) of a tear drop-shaped structure.

**Figure 3.**
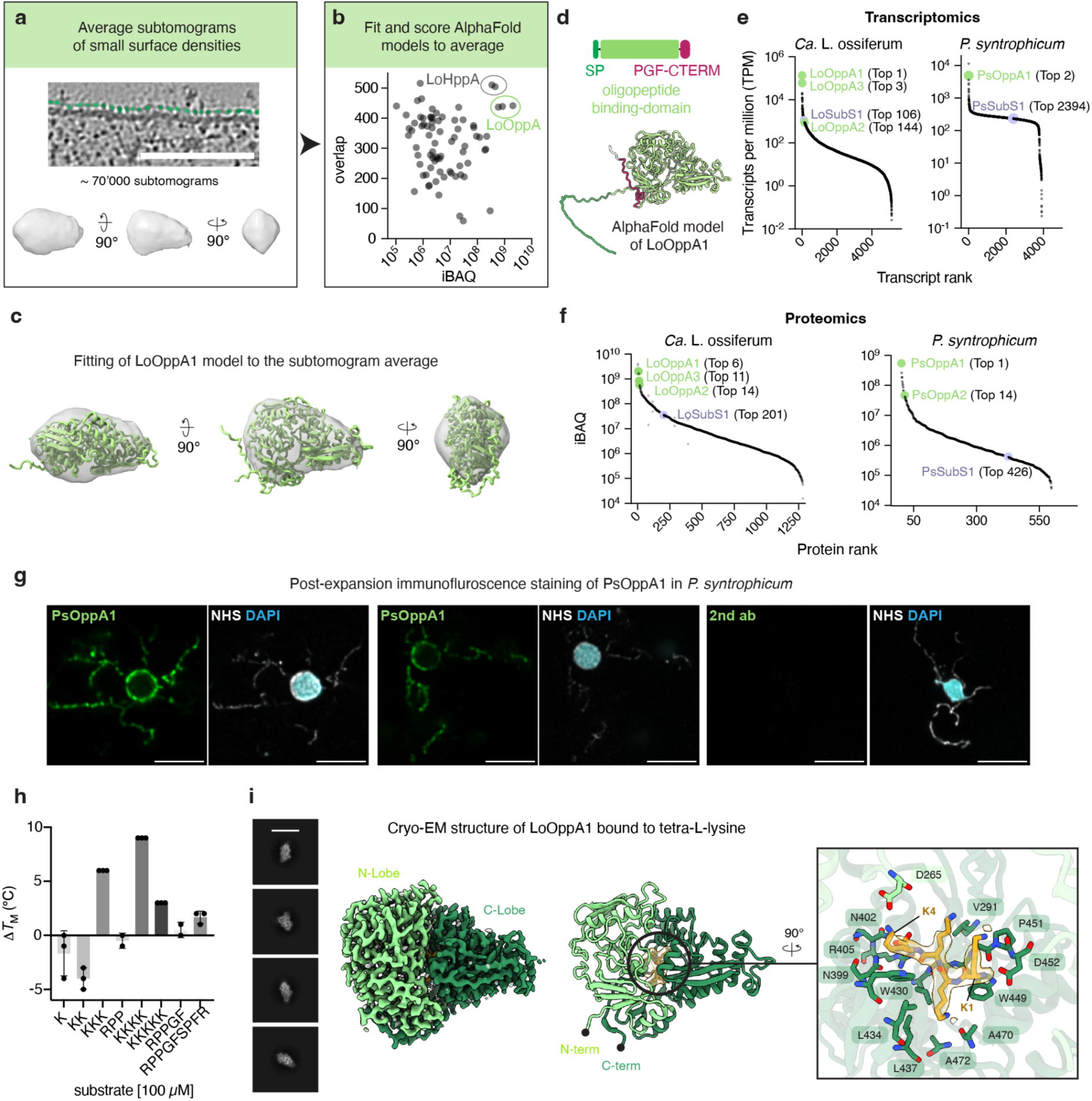
Oligopeptide-binding proteins are the major lokiarchaeal cell surface proteins. **a,** Visual surfaceomics workflow was applied to the SSDs of *Ca.* L. ossiferum. Shown are representative particles (top, in green) and the subtomogram average (bottom) at ∼18 Å resolution. Scale bar: 100 nm. **b,** Results of systematic fitting of all protein candidates show LoOppA homologs with a high overlap score and as the most abundant surface-bound proteins in the total proteome of *Ca.* L. ossiferum. See also Extended Data Fig. 5. **c,** Structural docking of the AlphaFold predicted structure of LoOppA1 to the reconstructed map. **d,** Domain organization of LoOppA1 based on both sequence and structure homology searches, showing the oligopeptide-binding region, a signal peptide (SP) and a PGF-CTERM domain commonly found in archaeal surface-anchored proteins. **e,** Gene expression analysis by transcriptomics of enrichment cultures of *P. syntrophicum* and *Ca.* L. ossiferum revealing OppA homologs among the most highly transcribed genes. Ranking of SubS1 transcripts are also indicated for comparison. **f,** Reanalysis of the whole cell proteomes of *P. syntrophicum*^40^ and *Ca.* L. ossiferum^16^ showing OppA homologs as the most abundant proteins detected from the total proteomes. **g,** Ultrastructure-expansion microscopy (U-ExM) of *P. syntrophicum* combined with NHS-ester bulk proteome labeling and post-expansion immunofluorescence staining using PsOppA1 primary antibodies and anti-rabbit IgG secondary nanobodies (2nd ab), revealing dense surface staining in both cell body and protrusions. Scale bars: 1 μm (biological scale). Shown are maximum intensity projections of central slices. Images on the right show a control experiment where the primary antibodies were omitted. The contrast of the secondary nanobody channel was adjusted equally between images. **h,** Shifts in the melting temperature (Δ*T*_M_) of purified LoOppA1 recorded through thermal shift assays upon addition of candidate substrates. The oligopeptide tetra-L-lysine (KKKK) induced the highest shift and was used as the ligand for cryo-EM single particle analysis. **i,** Single-particle cryo-EM structure of purified LoOppA1 incubated with 100 µM of tetra-L-lysine at 2.47 Å resolution. Shown are 2D classes (left), the density map and the corresponding atomic model colored according to the two lobes of the venus flytrap domain. A density consistent with the presence of tetra-L-lysine (orange) was observed in the substrate binding pocket of LoOppA1 and the residues participating in ligand binding are highlighted. Scale bar: 10 nm.

We next docked the previously generated structure library of predicted surface proteins into this reconstruction (Fig. 3b). Since the SSDs appeared to fully cover cell bodies and protrusions (Fig. 1), we expected the corresponding protein(s) to be highly abundant. We therefore plotted the candidates’ fitting scores against their abundance in the proteome, resulting in five top hits showing both high overlap scores and high abundance (Fig. 3b, Extended Data Fig. 5a). Two of these candidate proteins were excluded because they are membrane-integral sodium pumps lacking any predicted extracellular domains (Extended Data Fig. 5b). The remaining three candidates are homologous proteins, hereafter named LoOppA1/2/3 (LoOppA1, UYP46001.1; LoOppA2, UYP46002.1; LoOppA3, UYP44437.1), based on their annotation as oligopeptide-binding proteins (OppA; CDD COG0747, E-value [LoOppA1] = 1.05 × 10^−100^; PFAM PF00496, E-value = 3.6 × 10^−65^) (Fig. 3c,d, Extended Data Fig. 5a). OppA proteins are periplasmic or surface-bound proteins found in bacteria and archaea that primarily function in peptide scavenging from the environment, although they have also been found to be involved in other physiological processes^38,39^. Like LoSubS1, LoOppA1/2/3 contain N-terminal signal peptides and C-terminal PGF-CTERM domains (Fig. 3d, Extended Data Fig. 6), consistent with their translocation across the membrane and surface anchoring, respectively.

Interestingly, the genome of *Ca.* L. ossiferum encodes a total of 17 homologs of OppA (Extended Data Fig. 7a-c), seven of which were detected in the whole cell proteomics dataset (Extended Data Fig. 7b). LoOppA1/2/3 show highest expression, reflected in top rankings in the total transcriptome (Top 1, Top 144, Top 3, Fig. 3e) and proteome (Top 6, Top 14, and Top 11, respectively, out of 1’284 proteins, Fig. 3f). Notably, LoOppA1/2/3 were the most abundant membrane-bound proteins in the experimentally determined proteome (Extended Data Fig. 8a). Altogether, these findings suggest that the SSDs are predominantly composed of an assemblage of oligopeptide-binding proteins, dominated by LoOppA1/2/3.

Next, we asked whether OppA homologs played a similar role in *P. syntrophicum*, whose genome encodes four OppA homologs (Extended Data Fig. 9a). Reanalysis of *P. syntrophicum* whole cell proteomics data^40^ revealed that one of the OppA homologs, hereafter named PsOppA1 (QEE17258.1), constituted the most abundant protein in the total proteome (Fig. 3f), with its corresponding gene ranking Top 2 in the total transcriptome (Fig. 3e, Extended Data Fig. 9b). Like LoSubS1 and LoOppA1/2/3, PsOppA1 contains an N-terminal signal peptide and a C-terminal sorting domain (Extended Data Fig. 9c-e). Intriguingly, PsOppA1 was previously reported to be the most abundant glycoprotein in *P. syntrophicum* cultures^40^, similar to S-layer proteins in other archaea^40–43^. We therefore hypothesize that the SSDs found in *P. syntrophicum* tomograms are also predominantly composed of oligopeptide-binding proteins, dominated by PsOppA1.

To validate the localization and abundance of PsOppA1 on the surface of *P. syntrophicum* cells, we employed ultrastructure-expansion microscopy (U-ExM) with post-expansion immunofluorescence staining that we previously developed for *Ca.* L. ossiferum enrichment cultures^16^ (Fig. 3g, Extended Data Fig. 8b). To identify lokiarchaeal cells in the enrichment culture, we used N-hydroxysuccinimide (NHS) ester-dye conjugates that nonspecifically label total protein, allowing us to observe the distinct morphology of *P. syntrophicum*. Immunostaining with an antibody raised against a PsOppA1-specific peptide produced strong signals across the entire cell surface of all *P. syntrophicum* cells of different morphotypes (n = 171 cells across three cultures), consistent with extensive surface localization of PsOppA1 in *P. syntrophicum*.

### LoOppA1 binds oligopeptides and may form a complex with an ABC transporter

Next, we set out to test the putative oligopeptide-binding function of LoOppA1. To this end, we expressed it in *E. coli*, excluding the N- and C-terminal sorting domains (LoOppA1^34–564^), and purified it by immobilized metal affinity, anion exchange, and size exclusion chromatography (Extended Data Fig. 10a). Oligopeptide binding activity was then tested through a thermal shift assay^44,45^, in which an increase in melting temperature of ≥2 °C upon addition of a possible substrate was indicative of binding activity^46,47^. We tested mostly positively charged oligopeptides of different lengths. OppA homologs from other organisms favor such substrates because of negative surface charges within their substrate binding pockets^48,49^. LoOppA1 indeed showed a preference towards positively charged oligopeptides with lengths of three to nine amino acids (Fig. 3h). Tetra-L-lysine (KKKK) produced the largest thermal shift (Extended Data Fig. 10b).

In order to visualize ligand binding, we determined the LoOppA1 structure in the presence of tetra-L-lysine by single-particle cryo-electron microscopy (cryo-EM). The resulting structure, determined at 2.47 Å resolution, contained a density that is consistent with the presence of a tetra-L-lysine molecule in the expected substrate binding pocket of LoOppA1 (Fig. 3i, Extended Data Fig. 10c-j). In our structure, the two lobes of the venus flytrap domain are closed around the peptide substrate, closely resembling the closed conformations of previously determined oligopeptide-bound OppA structures from other organisms (Extended Data Fig. 11a)^49–51^. The specificity for oligopeptides is mediated by interactions between the N-terminal amine of the ligand and residues W449 and D452, and between the ligand C-terminal carboxylate and residues N399, N402, and R405. Extensive hydrogen-bonding interactions with the peptide backbone and limited specific side chain interactions, on the other hand, imply low sequence selectivity, as with other OppA proteins^39,51^. The electrostatic environment of the peptide-binding region nevertheless favors basic residues, while its large size could accommodate longer oligopeptides (Extended Data Fig. 11b,c). Interestingly, the predicted AlphaFold structure of LoOppA1 bound with tetra-L-lysine closely matched the experimentally determined structure (Cα RMSD = 0.886 Å, overall RMSD = 1.228 Å, Extended Data Fig. 11d). Altogether, these results demonstrate that LoOppA1 binds oligopeptides and suggest broad specificity with respect to oligopeptide length and/or sequence, consistent with a role in peptide uptake from the extracellular space.

Peptides captured by OppA proteins are typically delivered into the cytoplasm by an ATP-binding cassette (ABC) transporter complex. This complex comprises two integral membrane proteins OppB and OppC, which facilitate the passage of oligopeptides through the cell membrane, and two ATPases (OppD and OppF) that provide energy required to drive peptide translocation into the cytoplasm^52–55^. Gene clusters containing Lo*oppA3*, Ps*oppA1* and Ps*oppA2* contain the complete *opp* operon (Extended Data Fig. 12a). Notably, the predicted transporter components (OppBCDF) are also among the most abundant proteins detected in the proteome in both species, together with a subset of OppA homologs (Extended Data Fig. 12b). AlphaFold-Multimer predictions further support the assembly of the entire Opp transporter complex and a tetra-L-lysine ligand (Extended Data Fig. 12c). The high transcript and protein abundances of these components suggest a prominent role of peptide uptake in the physiology of both *Ca.* L. ossiferum and *P. syntrophicum*.

### Surface anchored subtilases and oligopeptide-binding proteins are conserved across Asgard archaea

Having identified the two major classes of surface proteins of two lokiarchaeal species, we next asked whether this surface architecture is conserved across Asgard archaea. We searched metagenomic datasets for surface-bound or secreted homologs of subtilase and OppA. Across 253 high quality Asgard archaeal genomes spanning all major phylogenetic lineages, we identified multiple homologs of both proteins, with an average of at least four homologs of subtilase and four homologs of OppA per genome (Extended Data Fig. 13a). This redundancy was particularly pronounced in Heimdallarchaea, where genomes contained an average of 7.8 SubS and 7.3 OppA homologs. Notably, the genome of the cultured hodarchaeon *Ca.* Margulisarchaeum peptidophila^12^ contains 24 homologs of SubS and 10 homologs of OppA. To determine whether these homologs found in other Asgard archaeal lineages have similar protein structures to that of lokiarchaeal subtilase and OppA, we performed structure similarity network analysis using AlphaFold-predicted structures (Fig. 4a,b). Despite stringent clustering criteria, most homologs grouped into a limited number of structural clusters, indicating broad structural conservation despite substantial sequence divergence (Extended Data Fig. 13b). Interestingly, the structures of the LoSubS1 and LoOppA1, which we identified as major cell surface proteins of *Ca.* L. ossiferum by visual surfaceomics, belong to large clusters encompassing homologs from diverse Asgard lineages.

**Figure 4.**
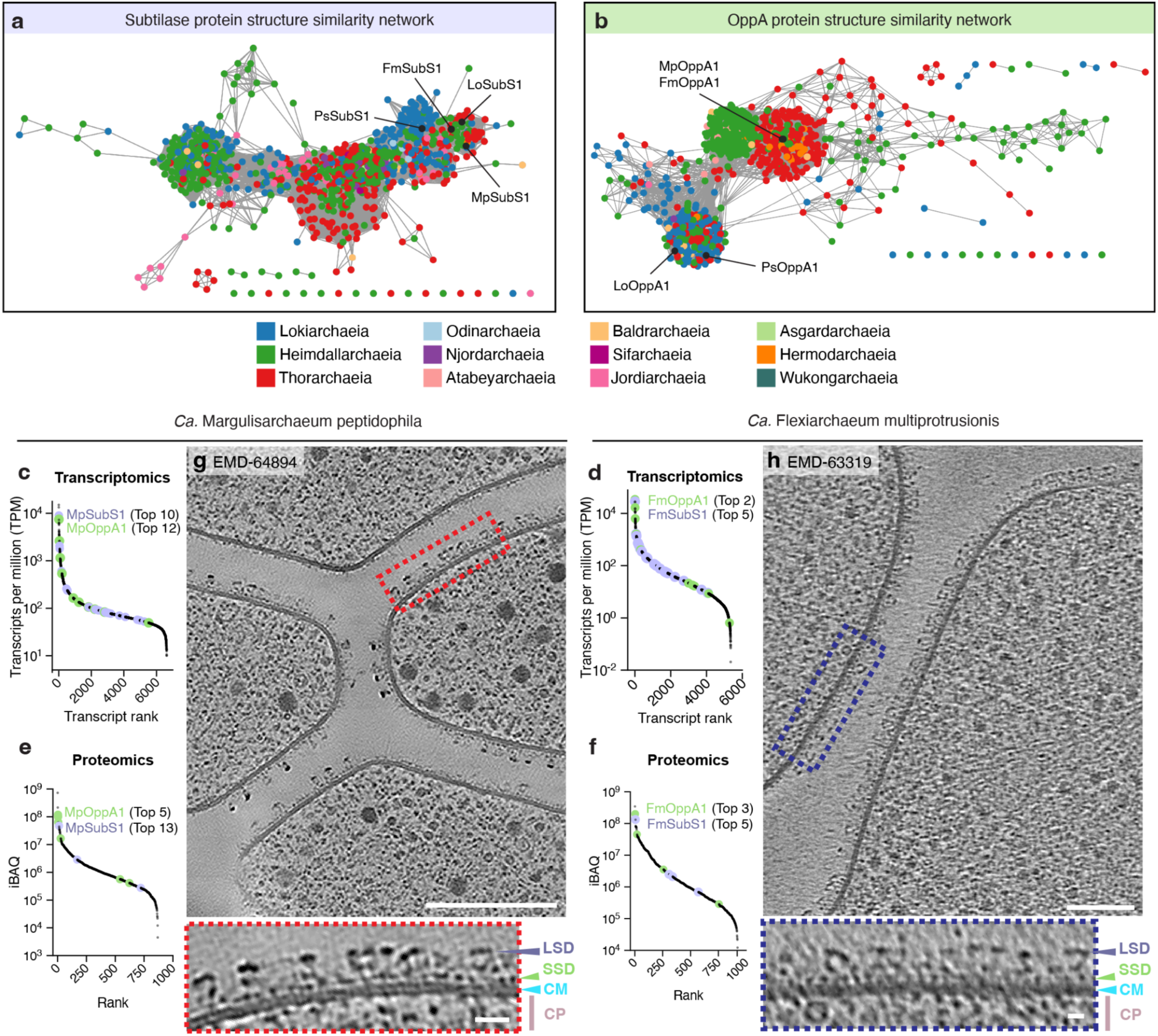
Surface anchored subtilases and oligopeptide-binding proteins are conserved across Asgard archaea. **a,b,** Protein structure similarity network of predicted surface-bound subtilase (a) and OppA (b) homologs identified in Asgard archaeal genomes, showing conserved structures across different lineages. The positions of selected homologs from different species are indicated. **c,d,** Reanalysis of transcriptomics-based gene expression profiling of the enrichment cultures of *Ca.* M. peptidophila (**e**) and *Ca.* F. multiprotrusionis (**f**) revealing that subtilase and OppA homologs are among the most transcribed genes for each strain^12^. **e,f,** Reanalysis of the whole cell proteomes of *Ca.* M. peptidophila^12^ (**g**) and *Ca.* F. multiprotrusionis^12^ (**h**) showing that subtilase and OppA homologs are among the most abundant proteins for each strain. **g,h,** Slices through cryo-tomograms of *Ca.* M. peptidophila (deposited previously as EMD-64894^12^, **i**) and *Ca.* F. multiprotrusionis (deposited previously as EMD-63319^12^, **j**) showing that their cell surface ultrastructure is similar to that of Lokiarchaea. Shown are also enlarged views of boxed areas, showing large and small surface densities (LSD/SSD) found on their cell membrane (CM). Scale bars: 100 nm and 10 nm for enlarged views.

Next, we set out to corroborate the relevance of subtilase and OppA homologs in recently reported Asgard archaeal enrichment cultures^12^. First, we assessed gene expression in *Ca.* M. peptidophila and *Ca.* Flexiarchaeum multiprotrusionis by reanalyzing transcriptomic and proteomic datasets^12^ (Fig. 4c-f). SubS and OppA homologs were among the most highly transcribed genes in both hodarchaeal strains, and the transcripts encoding the closest structural homologues of LoSubS1 and LoOppA1 were among the Top 10 transcripts, [hereafter named MpSubS1 (BEU39821.1) and MpOppA1 (BEU35157.1), and FmSubS1 (BFF48166.1) and FmOppA1 (BFF48142.1), for each strain respectively] (Fig. 4c,d). Proteomics data showed a similar pattern, with both homologs being among the most abundant proteins (Fig. 4e,f). Besides the analysis of gene expression levels, we also inspected previously deposited cryo-tomograms for both hodarchaeal species (EMD-64894 and EMD-63319). In both cases, the ultrastructure of the cell surface was similar to that of the previously observed lokiarchaeal species, featuring a cell membrane covered with two classes of small and large surface densities (Fig. 4g,h). Interestingly, in contrast to these lokiarchaeal strains, the surface of *Ca.* F. multiprotrusionis appeared more densely decorated with the large surface protein layer that putatively corresponds to FmSubS1 (Fig. 4h). This hypothesis is fully consistent with the proteomics data that indicated that FmSubS1 and FmOppA1 were expressed at comparable levels (Fig. 4f) - in contrast to *Ca.* L. ossiferum, where LoOppA1 expression was substantially higher compared to LoSubS1, resulting in a lower abundance of LSDs in cryo-tomograms.

Similarity analysis of predicted structures further showed that MpSubS1 and FmSubS1 share a very similar structure to LoSubS1 (Fig. 4a). Like LoSubS1, both proteins contain a globular protease domain connected to a stem composed of β-sandwich and Ig-like folds (Extended Data Fig. 14a). FmOppA1 and MpOppA1 differ from LoOppA1 by a small N-terminal extension, thereby placing them in another cluster in the network (Fig. 4b, Extended Data Fig. 14b). All of these homologous sequences contain an N-terminal signal peptide and C-terminal archaeosortase sorting signal (Extended Data Fig. 14c,d), consistent with surface localization. Taken together, these findings indicate that the cell surface composition we identified in lokiarchaeal strains is broadly conserved across other lineages of Asgard archaea.

### The cell surface is locally remodeled at contact sites

Finally, we asked whether this conserved cell surface architecture found in Asgard archaea may be locally remodeled to serve and adapt to different functions. We therefore cultured cells directly on EM grids and attempted to image *Ca.* L. ossiferum cells that were in contact with other *Ca.* L. ossiferum cells or in contact with other species. The obtained cryo-tomograms revealed a drastic change in the distribution of the putative subtilase densities, which were entirely absent at contact sites (intermembrane distance less than ∼25 nm). The distribution of the putative OppA densities appeared to remain unchanged in these regions. This pattern was observed for contact sites between *Ca.* L. ossiferum and *Ca.* L. ossiferum (n=1), between *Ca.*L. ossiferum and an intact bacterium (n=1), as well as between *Ca.* L. ossiferum and a lysing bacterium (n=2) (Fig. 5a,b, Extended Data Fig. 15). These observations are consistent with a spatial remodeling of the cell surface composition of *Ca.* L. ossiferum upon close contact with other cells.

**Figure 5.**
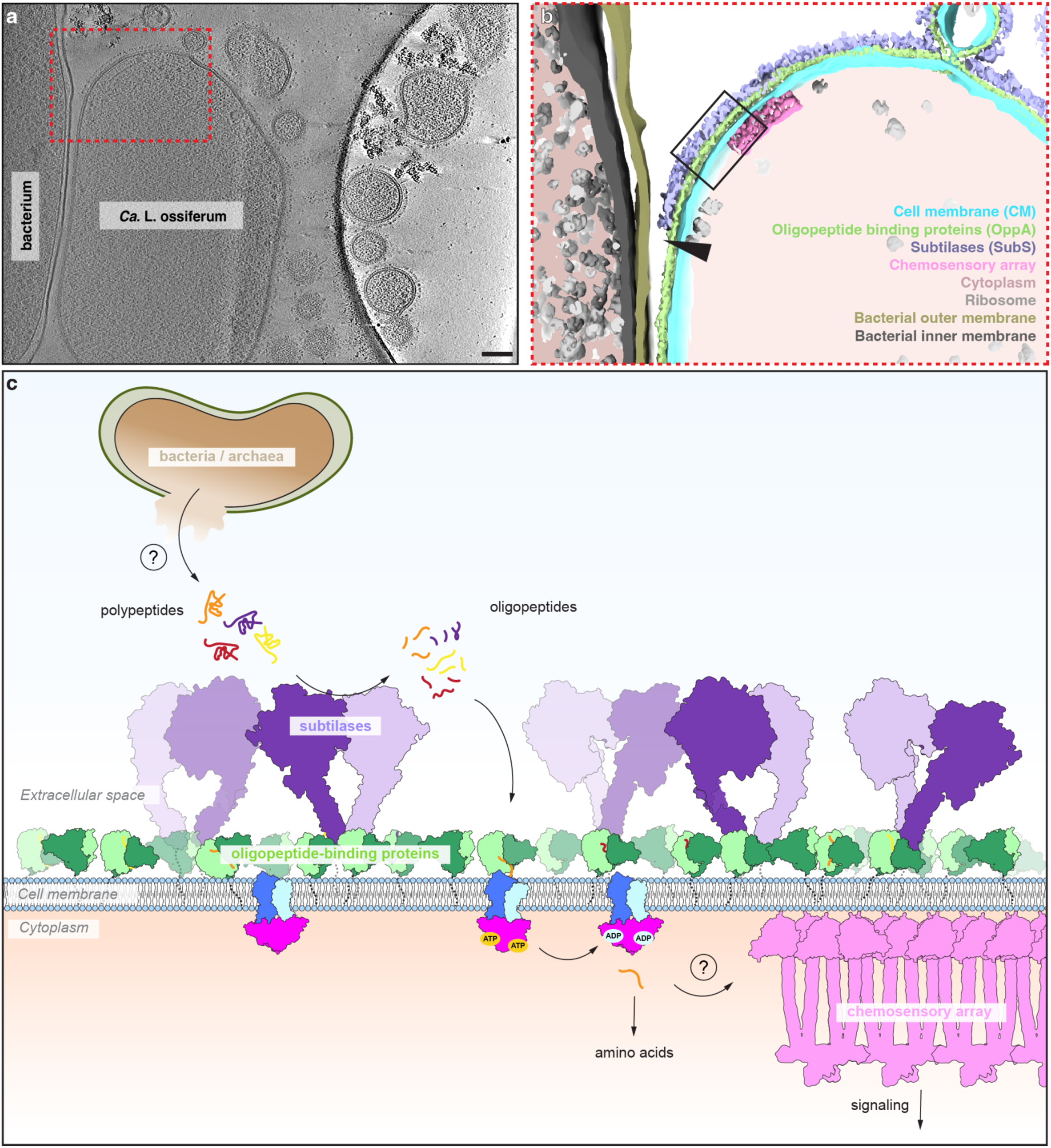
The flexible and substrate-scavenging surface of Asgard archaea. **a,b,** Slice through cryo-tomogram (**a**) and corresponding segmentation (**b**) of the observed cell-cell interaction site of a *Ca.* L. ossiferum cell in close association with an intact bacterium. Note the absence of LSDs and unchanged distribution of SSDs in regions of close membrane contact (indicated by black arrowhead), indicating a spatial remodeling of the *Ca.* L. ossiferum cell surface architecture. Scale bar: 100 nm. **c,** A schematic of the hypothesized flexible and catalytic cell surface of Asgard archaea. Membrane-bound subtilases cleave polypeptides from the extracellular space into smaller oligopeptides, which in turn are captured by surface-anchored oligopeptide-binding proteins. These oligopeptides are ultimately taken up by ABC transporters and may additionally be sensed by chemoreceptors.

## Discussion

### Visual surfaceomics

Cryo-electron tomography enables de novo identification of unknown structures in situ. At resolutions of 6-8 Å, subtomogram averages can often be docked with AlphaFold-predicted structures^56,57^. At lower resolutions, however, cross-correlation alone may be insufficient and often requires orthogonal multi-modal data and experimental validation^16,28,58^. This challenge was particularly pronounced for the lokiarchaeal cell surface, where slow growth, low biomass and non-axenic cultivation restrict conventional biochemical approaches, while the exceptionally small sizes of the target structures limit the obtainable resolution. To address this, we developed “visual surfaceomics” for identifying unknown surface-bound structures, allowing us to identify two of the smallest protein structures resolved to date by in-cell cryoET, demonstrating the potential of these approaches to extend the limits of structural characterization in situ. In the future, our methodological framework and similar recent approaches^28^ will enable the identification of further cell surface structures of Asgards and other (non-)model organisms.

### A substrate scavenging surface in Asgards

Visual surfaceomics identified two major protein classes on the lokiarchaeal cell surface: oligopeptide-binding proteins that densely cover the cell and more sparsely distributed subtilases (Fig. 5c). Comparative structural, genomic, transcriptomic, and proteomic analyses indicate that this surface protein composition and abundance is conserved across multiple Asgard archaeal lineages. This supports a model where surface-anchored subtilases cleave polypeptides into smaller oligopeptides, which are then captured by OppAs and shuttled into the cytoplasm via associated ABC transporters (Fig. 5c). This arrangement would spatially couple extracellular proteolysis to peptide uptake and provide a route for exploiting proteinaceous substrates from the environment. This unusual cell surface shows that peptide scavenging is central to the metabolic strategy of cultured Asgard archaea and agrees well with previous genomic analyses, which showed that Asgards lack several pathways for amino acid biosynthesis and are thought to scavenge them from the environment or acquire them from symbionts^1,59^. Moreover, the growth of cultured representatives (*P. syntrophicum, Ca.* M. peptidophila, *Ca.* F. multiprotrusionis) is dependent on peptides that cannot be replaced with a mixture of 20 individual amino acids^12^. Intriguingly, this specialized cell surface of Asgards may additionally be adapted towards the absorption of other kinds of substrates, aside from oligopeptides: proteomics analysis of hodarchaeal strains also showed high abundances of other predicted classes of substrate-binding proteins (in *Ca.* M. peptidophila, BEU36410.1 - sugar binding, Top 1; in *Ca.* F. multiprotrusionis, BFF51871.1 - nucleoside binding, Top 4; BFF48534.1 - Fe^3+^ binding, Top 7). These observations suggest that the Asgard cell surface likely represents a specialized metabolic interface that facilitates the uptake of diverse substrates from the surrounding environment.

### Multiple homologs for spatial remodeling?

Why do Asgard archaea encode and express multiple homologs of subtilases and OppAs? The first thought that comes to mind is that multiple copies may diversify the substrates that can be processed and acquired. A second possible reason for encoding multiple OppA homologs comes from our bioinformatic analysis. Intriguingly, some LoOppA homologs contain methyl-accepting chemotaxis protein (MCP) domains (LoOppA7 and LoOppA12, Extended Data Fig. 7). MCPs are chemoreceptors that sense different environmental cues through their ligand binding domains or indirectly through interactions with substrate binding proteins^60^, followed by signal transmission to the flagellum (bacteria) or archaellum (archaea)^61^. The involved proteins are typically organized in ordered cell envelope-associated chemoreceptor arrays^62^. In Asgards, putative chemoreceptor arrays have been observed previously^13^ and are seen in one of our cell-cell contact tomograms– oriented towards a bacterial cell (Fig. 5). Together with the report of actin-mediated motility in *Ca.* L. ossiferum and *Ca.* M. peptidophila^63^, OppA-MCP-mediated peptide sensing may enable directed motility towards a source of substrate. Third, carrying multiple homologs of cell surface proteins may introduce the possibility of spatial cell surface remodeling in order to serve distinct functions. The absence of subtilases at cell-cell contact sites (Fig. 5a,b, Extended Data Fig. 15) underlines that the general principle of localized remodeling is in fact implemented in the Asgard cell surface. Beyond the presence or absence of entire classes of cell surface proteins, it is further tempting to speculate that specialized OppA homologs may specifically localize to cell-cell contact sites and play a role in establishing and maintaining these interactions with other cells. While our data do not allow us to test this hypothesis at the moment, the report of some bacterial OppAs that play a role in adherence to cell surface glycosaminoglycans^64,65^ or glycoproteins^66^ of other organisms is consistent with this idea.

### Implications for eukaryogenesis

Finally, our findings also impact evolutionary hypotheses and agree with predictions made by the protrusion-mediated inside-out model of eukaryogenesis^9,10^. We confirmed the predicted absence of a paracrystalline S-layer that would allow the formation and remodelling of protrusions, likely mediated by eukaryotic signature proteins, such as actin and ESCRT^67–69^. This highly specialized cell envelope architecture, together with chemoreception (Fig. 5a,b), motility^63^, and spatial remodeling of the cell surface (Fig. 5a,b, Extended Data Fig. 15) may have been central to establishing and maintaining the close symbiotic interactions that led to the acquisition of the proto-mitochondrion.

## Data availability

Representative tomograms (EMD-100491, EMD-100493, EMD-100509, EMD-100510), subtomogram averages (SSDs, EMD-100522; LSDs, EMD-100523), and LoOppA1 cryoEM density map (EMD-100489) have been deposited in the EMDB. The LoOppA1 atomic model (PDB: 35FU) was deposited in the PDB.

## Author contributions

J.J.L.M. and F.W. developed the visual surfaceomics workflow and acquired, processed, and analyzed cryoET data. J.J.L.M., J.X., and L.D. performed subtomogram averaging of the LSDs and SSDs. J.J.L.M., J.X., J.C., and A.G. performed protein purification. D.A. acquired and analyzed cryoEM single particle data. J.J.L.M. performed biochemical assays and all bioinformatics analyses. F.W. and C.P.W. performed expansion microscopy and immunofluorescence experiments. F.W., C.P.W., A.K., and A.G. maintained *Promethearchaeum syntrophicum* cultures. F.W. and M.P. conceptualized and supervised the project. J.J.L.M., D.A., F.W., and M.P. wrote the manuscript with input from all the authors.

## Acknowledgments

We thank Christa Schleper and Thiago Rodrigues-Oliveira for providing cultures of *Ca.* Lokiarchaeum ossiferum. We thank Corina Hadjicharalambous, Tarun Iype, Maximilian Fottner (ETH Zürich), and Sihyun Sung (EMBL Grenoble) for helpful discussions with protein expression and purification; Pavel Afanasyev (ETH Zürich Cryo-EM Knowledge Hub) and Mary Dayne Tai (Centro Nacional de Biotecnología, Spain) for advice on cryo-EM sample preparation, data collection and processing; ScopeM and Yohei Yamauchi for instrument access at ETH Zürich; Paolo Nanni, Sibylle Pfammatter, and Jens Sobek (Functional Genomics Center Zurich) for proteomics and biophysics analysis; Fabian Eisenstein for advice on cryo-ET data collection and processing; Jürgen Jänes for support in running AlphaFold at scale; and Ariane Briegel (Institut Pasteur) and Adam Sidi Mabrouk (Leiden University) for helpful discussions on lokiarchaeal chemoreception. We acknowledge Hiroyuki Imachi and Masaru Nobu for providing access to *Ca.* M. peptidophila and *Ca.* F. multiprotrusionis proteomics data before publication. Calculations were carried out on the ETH Euler cluster. Pilhofer Lab members are acknowledged for helpful discussions. F.W. was supported by the European Molecular Biology Laboratory and an SNSF SPARK grant (229135). M.P. was supported by the European Research Council (101000232) and the NOMIS Foundation.

## Supplementary information

Supplementary Table 1. CryoEM data statistics of LoOppA1/tetra-L-lysine.

Supplementary Table 2. List of Asgard archaeal genomes used in this study.

Supplementary Table 3. List of predicted membrane-bound/secreted subtilase homologs identified from Asgard archaeal genomes.

Supplementary Table 4. List of predicted membrane-bound/secreted OppA homologs identified from Asgard archaeal genomes.

## Methods

### Culture conditions

*Prometheoarchaeum syntrophicum* MK-D1 was obtained as a co-culture with *Methanogenium sp.* MK-MG from Japan Collection of Microorganisms (JCM) as JCM 39240^T^. JCM 39240^T^ was cultivated in MKD1 medium (JCM medium #1339) supplemented with 0.05 % yeast extract as previously described^1,70^. *Ca.* L. ossiferum was grown as previously described^13^. Cultures were grown in sealed serum bottles filled with a head space gas mixture of 80:20 N_2_-CO_2_ at 20℃. Growth was monitored by routine qPCR as described^1,13^, and cultures with cell densities > 10^6^ mL were used for further investigation.

### Plunge freezing of *P. syntrophicum* and *Ca.* L. ossiferum co-cultures

Protein A-coated 10 nm gold beads (Cytodiagnostics) were mixed with aliquots of the culture at a 1:5 ratio and were kept at anaerobic conditions until plunge freezing. Copper EM grids (R2/1 or R2/2, Quantifoil) were glow-discharged, and around 3.5-4 µL of the cell suspension was applied. The grids were automatically blotted from the backside for 4-5 s and were plunged into liquid ethane/propane (37% (v/v) ethane)^71^ using an EM GP2 (Leica). For tomograms shown in Extended Data Fig. 15 and Fig. 5, *Ca.* L. ossiferum cultures were incubated on gold grids with a SiO_2_ mesh in a 96-well plate using a custom 3D-printed cell culture insert for 2 days prior to freezing in a Vitrobot Mark IV. All grids were finally stored in liquid nitrogen until imaging.

### Cryo-ET data collection

Tilt series were collected using a Titan Krios G4 FEG (Thermo Fisher Scientific) operated at 300 kV and equipped with a K3 camera and BioContinuum imaging filter (Gatan). SerialEM was used for data acquisition^72^. Owing to the low density of putative Asgard archaea cells, cryo-EM grids were first thoroughly screened at low magnification for target identification, as previously reported^13^. Tilt series were acquired in a dose-symmetric scheme that covers an angular range of −60° to +60° for a total electron dose of ∼160 e^−^/Å^2^. Tilt series were either acquired at nominal magnifications of 19,500 × (effective pixel size of 4.51 Å) with 2° increments and a defocus of −8 µm or 64,000 × (effective pixel size of 1.38 Å), acquired with 3° increments and a defocus values ranging from −3 to −7 µm. Tilt series used for subtomogram averaging were collected at 33,000 × (effective pixel size of 2.68 Å) acquired with 3° increments and defocus values ranging from −3 to −7 µm.

### Reconstruction, denoising, and segmentation of tomograms

Frame motion correction and summing of odd and even frames was performed using *alignframes* from IMOD v5.1.0^73^. Tilt series were aligned using gold fiducial markers, and tomograms were reconstructed at binning factors 1, 2, 4, 6, and 8 from the resulting tilt series using *etomo*^73^ or by patch tracking using AreTomo^74^. Tilt series alignments from *etomo* were also exported to WarpTools v2 for later subtomogram export and tilt series refinement^75,76^. Models for denoising and missing wedge reconstruction were trained and applied using DeepDeWedge v1^77^ or IsoNet v0.3^78^. Denoised tomograms were then segmented using Dragonfly v2022.2 (Comet Technologies Canada) as previously described^19^. A training dataset sized 79 x 75 x 63 pixels from a tomogram (Extended Data Fig. 1) was manually segmented with four classes (cell membrane, LSDs, SSDs, and background) and was used for training a U-Net model (input dimension = 2.5D, input slices count = 5 slices). The model was reiteratively improved by adding more manually segmented slices. Membrane segmentation was improved by several morphological operations (dilate and close) to achieve segmentations with no holes. The resulting model was applied to tomograms belonging to the same dataset, and were used for particle picking of SSDs. New models were likewise prepared for segmentation of other tomograms (Fig. 1a-d, Fig. 5a,b, Extended Data. Fig. 15). Segmentations were visualized with ChimeraX v1.9^79^.

### Particle picking of small and large surface densities from *Ca.* L. ossiferum

Models for particle selection of LSDs from denoised tomograms (binning factor of 6) were trained and applied using crYOLO v1.9.9^20^. Particle coordinates for the SSDs were generated from Dragonfly segmentations (binning factor of 4) as previously described^19^. Briefly, particles were first separated using a watershed transform and rounds of morphological operations (erode). All particle coordinates were generated from weighted centers of masses in x, y, and z (centroids). Particle tables were then converted to Dynamo table files (*.*tbl*) for downstream processing. Misannotated particles were manually removed using the ArtiaX v0.3 plugin in ChimeraX^80^ or boxmanager from napari v0.4.17^81^. Initial orientations for the particles of SSDs were imparted based on membrane geometry through *dpktbl.triangulation.fillTable* in Dynamo v1.1.532^82,83^ using membrane meshes derived from Dragonfly segmentations (Extended Data Fig. 4a). Membrane segmentations from Dragonfly were converted to membrane meshes using Chimera^84^.

### Geodesic nearest-neighbor distance and particle density analysis

Geodesic nearest neighbors and particle densities were computed using trimesh^85^ and potpourri3d^86^ as previously described^87^ with modifications. Only particles with a maximum shortest Euclidean projection distance to the mesh of 40 nm and 10 nm were used in the analysis for LSDs and SSDs, respectively. Geodesic distances were then calculated using the heat method^88^.

### Subtomogram averaging of LSDs from *Ca.* L. ossiferum

Initial subtomogram averaging of the LSDs was performed in Dynamo^82^. Particles were cropped from DeepDeWedge-denoised tomograms at a binning factor of 6 (pixel size of 16.07 Å) and box size of 48^3^ (52,930 particles) using *dtcrop*. The azimuth angles were randomized through *dynamo_table_randomize_azimuth* and were used for coarse alignment of the particles in *dcp*. The particles with refined orientations were cropped using WarpTools from unfiltered tomograms at a binning factor of 4 (pixel size of 10.71 Å) and box size 36^3^. Iterative refinement was then performed in RELION using the reconstruction obtained from *dcp* as initial reference. The refined particles were then subjected to two rounds of 3D classification without angular sampling (K=5, T=0.5) and refinement to remove bad particles. The filtered particles (39,162 particles) were then recropped at a binning factor of 2 (pixel size of 5.36 Å) and box size 72^3^ using WarpTools and were refined in RELION. A final reconstruction of the LSDs at an estimated resolution of ∼15 Å was determined from these subtomograms without applying symmetry (Extended Data Fig. 2).

### Subtomogram averaging of SSDs from *Ca.* L. ossiferum

Subtomogram averaging of the SSDs was performed in Dynamo^82^. Particles with initial orientations were cropped from IsoNet-denoised tomograms at a binning factor of 4 (pixel size of 10.71 Å) and box size of 24^3^ (73,654 particles) using *dtcrop*. The azimuth angles were likewise randomized through *dynamo_table_randomize_azimuth* and were used for coarse alignment of the particles in *dcp*. 2D projections of each refined particle was calculated as previously described^13,16^ and were subjected to 2D classification with angular sampling (K=30, T=1) using RELION. Particles with a clear membrane signal were then selected for downstream refinement in Dynamo. Filtered particles (46,446 particles) were recropped from unfiltered tomograms with *dtcrop* at unbinned pixel size (2.68 Å) with a box size of 96^3^. The particles were then split into half-datasets using *dteo* and were iteratively refined using *dcp* starting at a binning factor of 4 (pixel size of 10.71 Å) and eventually at binning factor of 2 (pixel size of 5.36 Å). Resolution was calculated using *dfsc* based on the Fourier shell correlation^89^ of the reconstructions from both half-datasets. The final reconstruction of the SSDs was determined at an estimated resolution of ∼19 Å (Extended Data Fig. 4b).

### Analysis of transcriptomics and proteomics data

Raw RNA-seq reads were obtained from Sequence Read Archive (NCBI-SRA) of runs SRR32148623 for *Ca.* L. ossiferum^16^, DRR199588 for *P. syntrophicum*^1^, DRR620298 for *Ca.* M. peptidophila, and DRR620299 for *Ca.* F. multiprotrusionis^12^. RNA-seq data was analyzed as previously described, with modifications^16^. Adapter trimming and low quality read filtering was performed with fastp v0.24.1^90^, and rRNA reads were removed using SortMeRNA v4.3.6^91^. Filtered, high quality reads were annotated against their respective genomes and sorted using bowtie2 v2.5.4^92^. Quantification and normalization into transcripts per million (TPM) was performed StringTie v3^93^. Data was ranked by TPM value separately for each organism and was plotted as TPM value vs. transcript rank using GraphPad Prism v10.6.1. Proteomics data of *Ca.* L. ossiferum (PRIDE: PXD-059733) was previously analyzed and reported^16^. *P. syntrophicum* (PXD-051006)^40^, *Ca.* M. peptidophila (PXD-059994)^12^, and *Ca.*F. multiprotrusionis (PXD-059995)^12^ was re-analyzed as previously determined^16,94^. Briefly, raw MS data was processed by MaxQuant v2.0.10 and proteins were identified through the integrated Andromeda search engine to produce iBAQ values^95^. A false discovery rate (FDR) of < 0.1% was used to filter peptide identifications, while an FDR of < 1% was used to filter protein identifications, given that it contains at least 2 identified peptides. Detected proteins were ranked based on their iBAQ values to determine protein rank.

### AlphaFold structure prediction of putative surface-bound proteins of *Ca.* L. ossiferum

The annotated complete genome of *Ca.* L. ossiferum (GCA_025839675.1) was obtained from NCBI. PFAM domains were identified from all protein sequences using *hmmscan* from HMMER v3.3.2^96^ with the Pfam HMM database v35.0^97^. Subcellular localization was predicted through DeepTMHMM^98^ and all proteins annotated to contain either a signal peptide (SP), a transmembrane region (TM), or both (SP+TM) were considered as putative membrane bound or secreted proteins. From these filtered proteins, only those which are detected from proteomics analysis were used for AlphaFold v2.0.1-based protein structure prediction (152 proteins) using default parameters^30^. Only the top ranked model was used for subsequent analysis. AlphaFold v3^99^ was used to predict the structures of MpSubtS1, MpOppA1, FmSubS1, FmOppA1, and multimeric structures of OppABCDF and tetra-L-lysine.

### Fitting of AlphaFold predictions to surface protein reconstructions from *Ca.* L. ossiferum

Fit libraries representing the best possible positions of each predicted AlphaFold structure into each of the calculated reconstructions were calculated using the Efitter pipeline in Assembline v1.05^31^. Briefly, predicted structures were fitted against the reconstructed maps at a resolution of 21.4 Å for the SSDs and 10.71 Å for the LSDs using 10,000 initial random placements. For the LSD, the map corresponding to the putative monomer was used for the fitting. Fits with the best overlap value were used and were plotted as overlap value vs detected iBAQ value for each protein.

### Antibody generation

Affinity purified polyclonal antibodies from rabbits were generated by Eurogentec. Specific peptides against PsOppA1 (83-97: CLLAKEMGNWSADNLN, 359-373: CFLIDDQSELENPVPA) were used for antibody generation. Antibodies were then validated by Western blotting. From the enrichment culture, 750 µL was used per well by harvesting cells through centrifugation at 20,000 × g for 30 min at 4°C. Cell pellet was directly lysed in SDS loading buffer (Bio-Rad) supplemented with 1% (v/v) β-mercaptoethanol at 95 °C. Cell lysates were run on 4–20% Tris-Glycine gradient gels (Bio-Rad) and were transferred onto nitrocellulose membranes. After blocking with 5% (w/v) non-fat dry milk in TBST, the membranes were probed with each PsOppA1-specfic primary antibody and secondary (goat anti-rabbit HRP, Invitrogen, 31460) antibody. Chemiluminescent signals were detected using enhanced chemiluminescence (ECL) substrate.

### Ultrastructure Expansion Microscopy (U-ExM)

U-ExM analysis of PsOppA1 was done essentially as described^16,100^. *P. syntrophicum* cultures were fixed in formaldehyde/glutaraldehyde (4/0.025 % [w/v]) for 20 min, immobilized on PEI-coated coverslips and incubated in anchoring solution (2% acrylamide and 1.4% formaldehyde in PBS) for 3 h at 37 °C before gelation in monomer solution (19% sodium acrylate, 10% acrylamide and 0.1% N,Nʹ-methylenbisacrylamide in PBS) supplemented with 0.5% tetramethylethylendiamine and ammonium persulfate for 1 h at 37 °C. Gels were denatured at 85 °C for 1.5 h in denaturation buffer (200 mM SDS, 200 mM NaCl and 50 mM Tris-HCl, pH 9) and fully expanded in ddH_2_O. For immunostaining with PsOppA1 antibodies, gels were washed in PBS and subsequently incubated for 3 h at 37 °C with antibodies against PsOppA1 (diluted 1:100 in MaxBind), washed three times with MaxWash and incubated with an Atto643-labeled anti-rabbit nanobody (NanoTag N2404-At643-S, 1:200 diluted in MaxBind) overnight at 4 °C. After washes in MaxWash and counterstaining with NHS-Atto488 (4 μg/ml) and DAPI (2 μg/ml) in PBS for 1 h, gels were fully expanded in ddH_2_O and z-stacks were acquired using a Zeiss LSM990 Airyscan microscope and a 63x 1.4 NA oil immersion objective. Images were processed using Airyscan joined deconvolution (Fig. 3g) or Airyscan 3D Processing (Extended Data Fig. 8b) in Zen Blue and central slices were extracted in Fiji^101^.

### LoOppA1 overexpression and purification

Codon optimized LoOppA1 (UYP46001.1, without N-terminal signal peptide (amino acids 2-33) and C-terminal transmembrane domain (amino acids 565-590)) with a C-terminal tobacco etch virus (TEV) protease cleavage site and His_6_-tag was synthesized and cloned into vector pET28a by Genscript. This construct was transformed to *E. coli* BL21 (DE3) for protein overexpression. From an overnight culture, a 1% protein overexpression culture was prepared in 2X YT medium and was incubated at 37 °C with shaking at 220 rpm. After growth to an OD of 0.7, the culture was cooled and 0.5 mM IPTG was added. The culture was further incubated at 16 °C with shaking at 180 rpm overnight. The cells were then harvested by centrifugation at 10 000 rpm for 10 min at 4 °C. The cells were resuspended in lysis buffer (25 mM HEPES, 500 mM NaCl, 1 mM DTT, 50 ug/mL DNAse (Roche), 1 × protease inhibitor cocktail (Roche), 40 mM imidazole, 0.1% yeast extract, 0.1% tryptone, pH 7.4) and were lysed on ice using sonication for 5 minutes total of 10 s on/20 s off at 50% power. Cell debris was separated by centrifugation at 20 000 rpm for 30 min at 4 °C and passing through a 0.22 µm syringe filter.

Protein purification was performed on an ÄKTA pure chromatography system. Overexpressed LoOppA1 was initially purified from the cell lysate through immobilized metal affinity chromatography by using a HisTrap FF column (Cytiva), which was equilibrated with the wash buffer (25 mM HEPES, 1 M NaCl, 1 mM DTT, 4 mM ATP, 20 mM MgCl_2_, pH 7.4). After several washing steps with the wash buffer, the proteins were eluted with a linear gradient (0-100%) of elution buffer (25 mM HEPES, 500 mM NaCl, 500 mM imidazole, pH 7.4). Fractions containing overexpressed protein were further purified through ion exchange chromatography by using a HiTrap Capto Q ImpRes column (Cytiva) equilibrated with a buffer containing 25 mM HEPES pH 7.4. A linear gradient (0-100%) of a buffer containing 25 mM HEPES, 1 M NaCl, pH 7.4 was used to elute LoOppA1. A polishing step was performed through gel filtration chromatography on a Superdex 200 Increase column (Cytiva) pre-equilibrated with a buffer containing 25 mM HEPES, 500 mM NaCl, pH 7.4. Purity of fractions were investigated using SDS-PAGE and purified LoOppA1 fractions were combined and concentrated using a 50 kDa centrifugal filter (Merck) with the addition of 10% glycerol. Protein concentration was estimated using Nanodrop (Thermo Fisher) and aliquots were snap-frozen using liquid nitrogen and stored at −80 °C.

### Thermal shift assay

Reaction mixtures were performed in duplicate or triplicate and were prepared in clear 96-well PCR plates. Purified LoOppA1 was diluted to 2.5 µM in assay buffer (25 mM HEPES, 500 mM NaCl, pH 7.4), and for conditions with ligands, 100 µM of ligand was added in a total reaction volume of 22.5 µL. Reactions were incubated for 30 minutes at room temperature, after which 5 × SYPRO Orange Protein Gel Stain was added to a total reaction volume of 25 µL. The plates were sealed with an optically clear sealing film and were centrifuged at 1 000 g for 1 min. The plates were placed on a Bio-Rad CFX96 Touch Real-Time PCR System in which the temperature was ramped from 25 °C to 100 °C at a rate of 1 °C/min. Fluorescence was measured using the FRET channel. Melt curve analysis and melting temperature determination was performed using Bio-Rad CFX Maestro.

### Sample preparation for single-particle Cryo-EM

Purified LoOppA1 at a concentration of 1 mg/mL in assay buffer containing 25 mM HEPES, 200 mM NaCl, pH 7.4 was incubated with 100 µM of tetra-L-lysine at room temperature for 30 minutes. Shortly before freezing, octyl glucoside (Anatrace) was added to the sample at a final concentration of 0.05% w/v. 4 µL of sample was then applied to freshly glow-discharged (10mA, 60s) UltraAuFoil (Quantifoil) grids in a Leica EM GP2 plunge freezer (Leica Microsystems) and blotted for 2-4 s before plunge freezing in a liquid ethane-propane mixture.

### Cryo-EM single-particle data collection

Details of cryo-EM data collection are summarized in Supplementary Table 1. Micrographs were collected using a Titan Krios (Thermo Fisher Scientific) operated at 300 kV and equipped with a K3 direct electron detector operated in CDS mode and a BioContinuum imaging filter (Gatan). EPU was used for data collection. All movie stacks were recorded in counting mode with 40 frames per stack and a total electron dose of ∼53 e-/Å^2^. A nominal magnification of 165,000 × (effective pixel size of 0.51 Å), total exposure time of 3 s, and defocus values ranging from −0.4 to −1.8 µm were used during data acquisition.

### Cryo-EM image processing and model building

A total of 12,032 micrographs were imported into CryoSPARC v5 beta and motion corrected using Patch motion correction. The micrographs were Fourier cropped to a pixel size of 1.02 Å/px and CTF parameters were estimated with Patch CTF. Micrographs with an estimated CTF fit worse than 7 Å were excluded, resulting in a curated dataset of 11,035 micrographs. These micrographs were then denoised using the Micrograph denoiser job and particles were picked using the Blob picker with an elliptical template. Particles picked on the gold foil or in bad ice were excluded using the Micrograph junk detector job and good particles were subsequently extracted with a box size of 240px, Fourier cropped to 120px. After one round of 2D classification (200 classes, max. resolution 3 Å, circular mask diameter 110 Å, 80 online-EM iterations, batch size per class 400), classes showing the characteristic teardrop shape of OppA and unambiguous high-resolution features were selected, resulting in 1,173,041 particles. One round of Ab-initio reconstruction (3 classes, 600,000 particles, initial resolution 12 Å, max resolution 9 Å, initial minibatch size 300, final minibatch size 1000) and two rounds of heterogeneous refinement (3 classes, initial resolution 10 Å) resulted in one good class containing 500,541 particles. The particles were re-extracted at the full box size of 240px and non-uniform refinement was applied. Further processing with reference-based motion correction and extraction at the original pixel size of 0.51 Å/px, followed by several iterations of CTF refinement and non-uniform refinement finally yielded a map at a nominal resolution of 2.47 Å, calculated from 498,446 particles. The default CryoSPARC parameters were used unless otherwise indicated.

The predicted AlphaFold3 model of LoOppA1 with bound tetra-L-lysine was docked into the final map via rigid body fitting in ChimeraX and iteratively refined using Coot and *phenix.real_space_refine*. *Phenix.molprobity* was then used to validate the model. Validation statistics are summarized in Supplementary Table 1. All figures were prepared in ChimeraX.

### Comparative genomics of Asgard archaea

High quality genomes of Asgard archaea were identified from the Genome Taxonomy Database (GTDB) Release 10-RS226 (16^th^ April 2025)^102^ with the criterion “GTDB species representatives only”. After filtering, 251 genomes were downloaded using NCBI Datasets^103^ and their GTDB taxonomic assignment was used during further analysis. Additionally, genomes of *Ca.* M. peptidophila HC1 (AP029002.1) and *Ca.* F. multiprotrusionis (AP029247.1) were included in the analysis^12^. Protein sequences were reidentified from all genomes using Prokka v1.14.5^104^ for standardization. Homologs of OppA and SubS were identified by running *hmmscan* on all proteins against Pfam family HMMs PF00496.27 for OppA and PF00082.27 for Subt. Only hits with a domain bitscore greater than the thresholds for each HMM (32.4 for OppA and 21.8 for Subt) were considered. For all identified OppA and SubS homologs, DeepTMHMM was used to filter homologs that are putatively membrane-bound or secreted proteins. Putative OppA homologs were also searched against the Transporter Classification Database (TCDB)^105^ to filter out sugar-binding and nickel-binding proteins. *opp* gene clusters were identified through the individual Pfam domains of each gene. Gene clusters were visualized using *clinker* from CAGECAT v3.1^106^.

### Structure-based similarity networking of OppA and SubS homologs

Protein structures of all identified putatively membrane bound/secreted SubS and OppA homologs were predicted using AlphaFold v3^99^. TM-scores^107^ between predicted protein structures were calculated using FoldSeek v10-941cd33^108^ easy-search in all vs all format. Networks were built using Cytoscape v3.10.4^109^ with edge distance calculated as *D = 1 – TM-score*, and visualized using a Prefuse Force Directed OpenCL layout. Only protein pairs with a TM-score > 0.5 and with more than 80% alignment coverage for both query and target proteins were retained in the network. Percent identity of amino acid sequences between proteins were calculated using MMseqs2^110^ easy-search.

---

**Extended Data Fig. 1.**
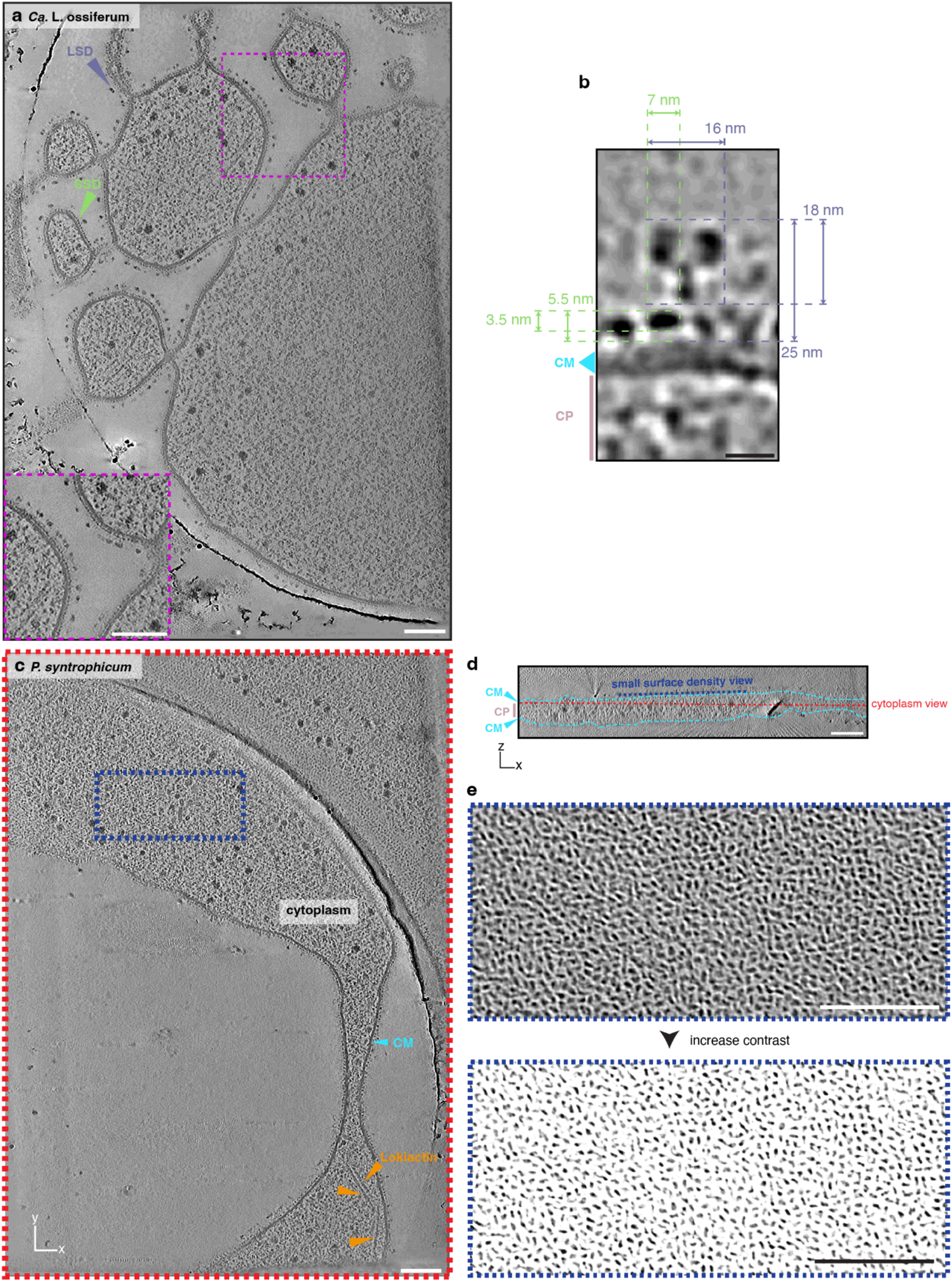
Distribution of surface densities in cell bodies and protrusions of Lokiarchaea. **a,** Additional cryo-tomographic slice through a *Ca.* L. ossiferum cell revealing small surface densities (SSD) and large surface densities (LSD). Inset shows an example of high LSD density (see also Fig. 1a). Scale bars: 100 nm. **b,** Dimensions of the small and large surface densities of *Ca.* L. ossiferum. Same as Fig. 1A inset. Scale bar: 10 nm. **c,d,** Cryo-tomographic slices of *P. syntrophicum* shown in top view (xy plane; **c**) and perpendicular view (xz plane; **d**). The z-height of the slice shown in (**c**) is indicated in (**d**). The blue dashed box indicates the field of view shown in **e**. Putative Lokiactin filaments indicated by orange arrowheads. Scale bars: 100 nm. **e,** Top view of the *P. syntrophicum* cell surface (field of view is indicated in panel **c**; z-plane is indicated in panel **d**) shows the unordered and dense arrangement of the SSDs. Scale bars: 100 nm.

**Extended Data Fig. 2.**
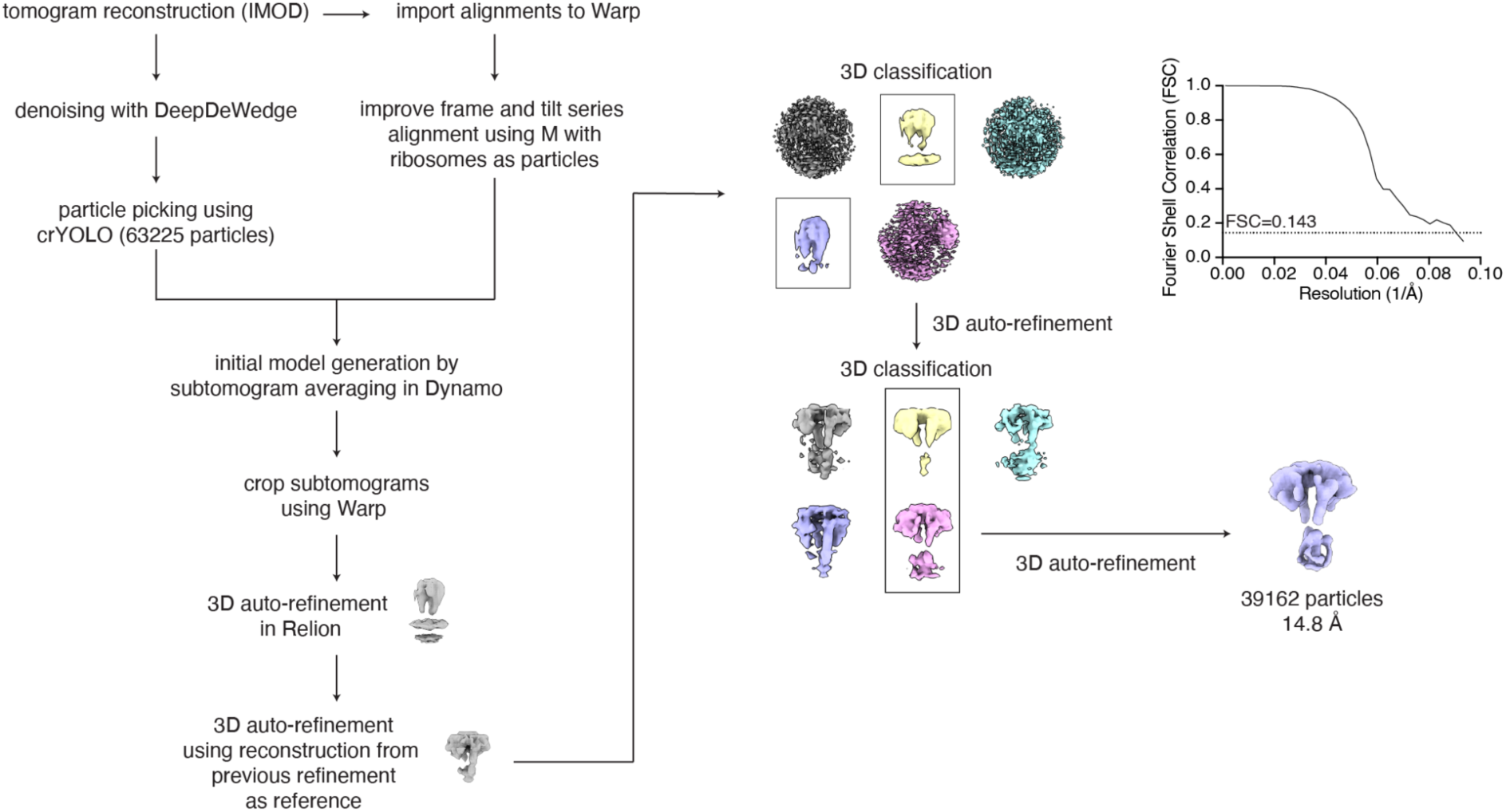
Subtomogram averaging of the LSDs from *Ca.* L. ossiferum. Data processing flowchart for the particle picking of the LSDs and subsequent subtomogram averaging. For details, see Materials and methods.

**Extended Data Fig. 3.**
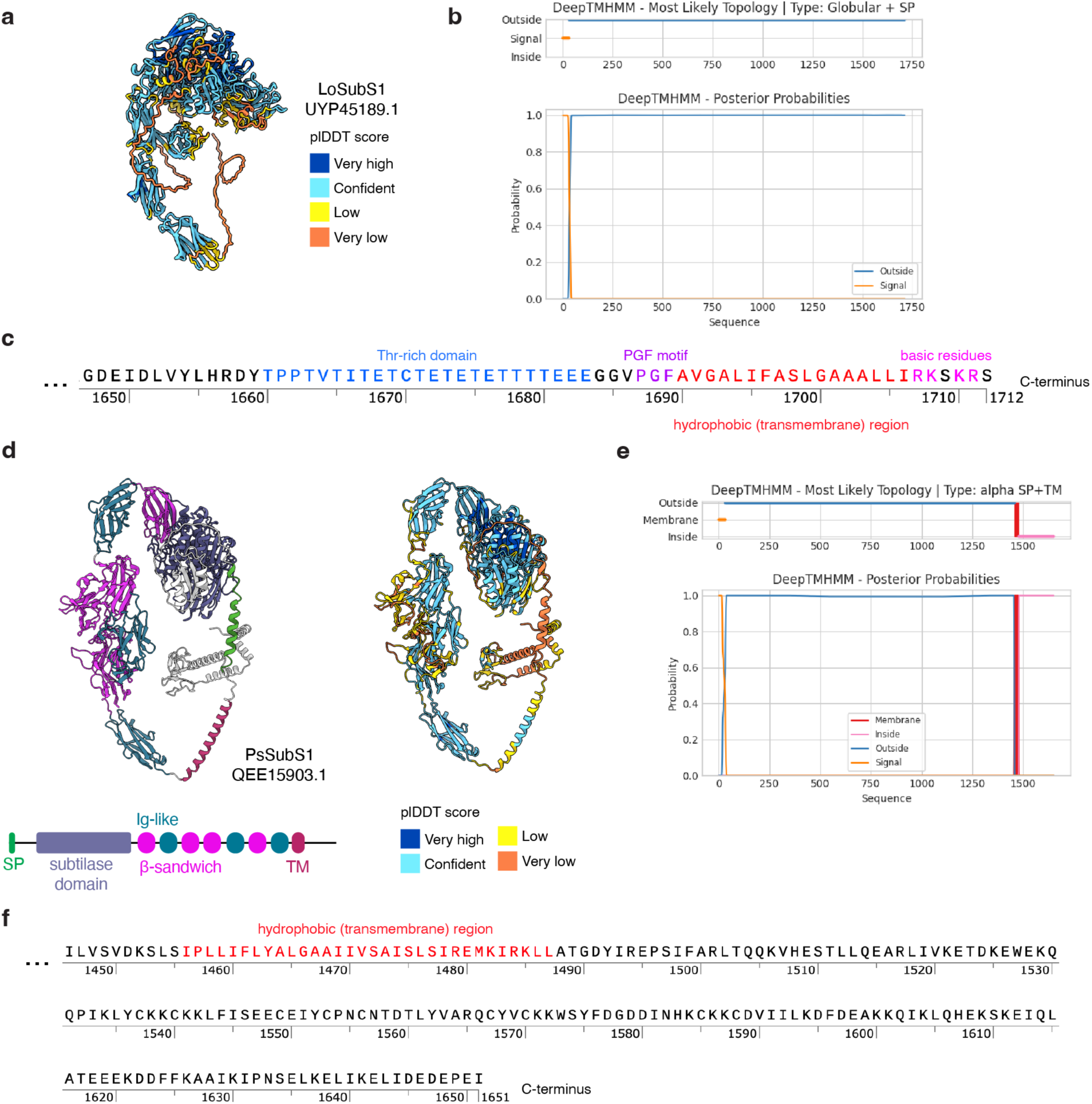
Structure and topology prediction of LoSubS1 and PsSubS1 and their sequence features. **a,** AlphaFold predicted structure of LoSubS1, colored by plDDT score. A pTM value of 0.74 is calculated from the model, indicating a high quality and confident prediction. **b,** Protein topology prediction of LoSubS1 using DeepTMHMM. LoSubS1 is predicted to be a secreted globular protein containing an N-terminal signal peptide. **c,** Sequence features of the PGF-CTERM sorting signal domain found in the C-terminus of LoSubS1. **d,** Predicted structure of PsSubS1 colored by structural domains (left) and plDDT score (right). Note the similar domain organization to that of LoSubS1. SP, signal peptide; TM, transmembrane region. **e,** Protein topology prediction of PsSubS1 using DeepTMHMM. PsSubS1 is predicted to be a secreted globular protein containing an N-terminal signal peptide with a transmembrane domain. **f,** Transmembrane region that extends to a cytoplasmic region found in the C-terminus of PsSubS1.

**Extended Data Fig. 4.**
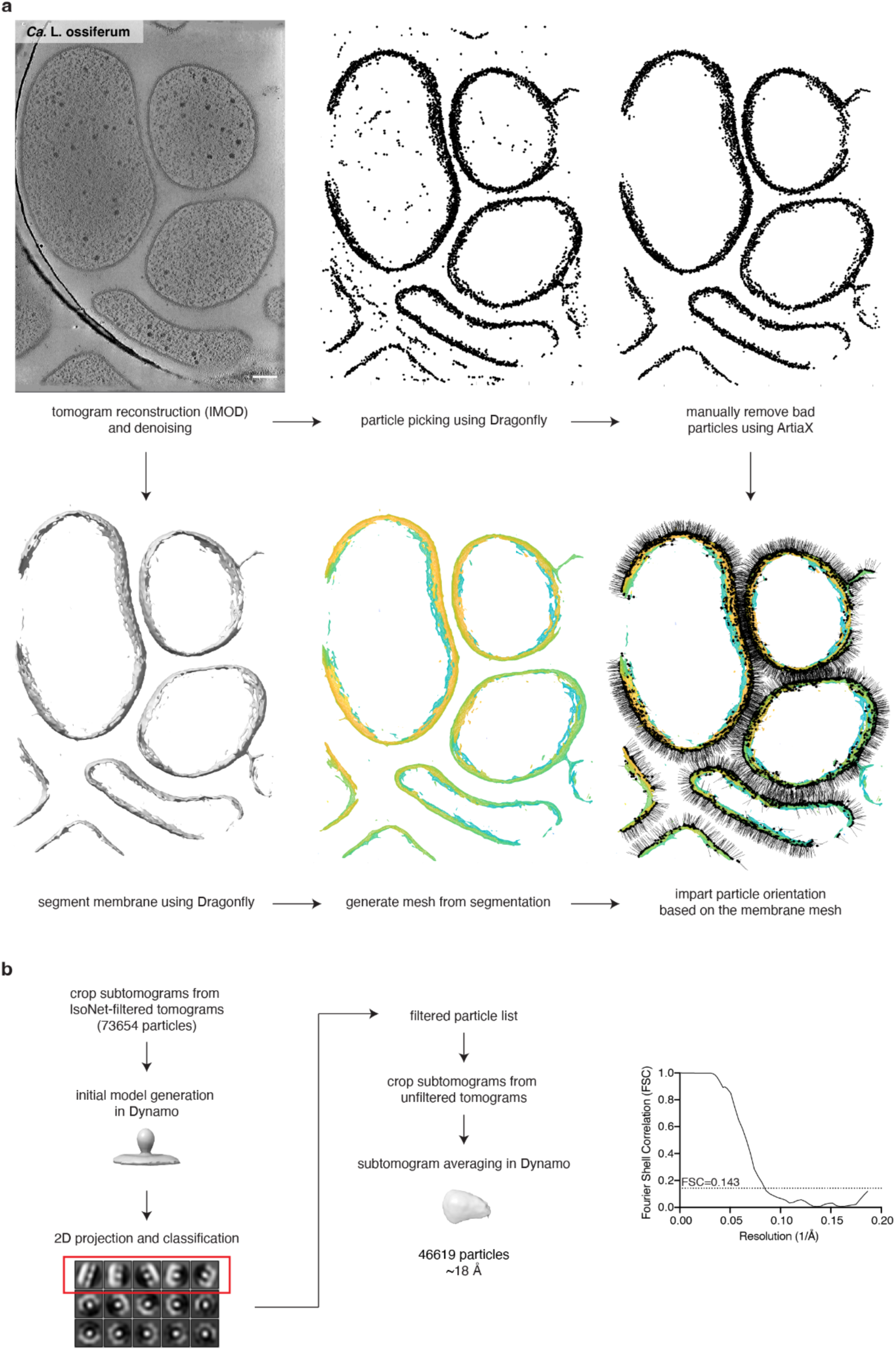
Particle picking and subtomogram averaging of the SSDs from *Ca.* L. ossiferum. **a,** Data processing flowchart for the curation of SSD particle coordinates for subtomogram cropping and imparting initial particle orientations. Scale bar: 100 nm. **b,** Data processing flowchart for the subtomogram averaging of SSDs. For details, see Materials and methods.

**Extended Data Fig. 5.**
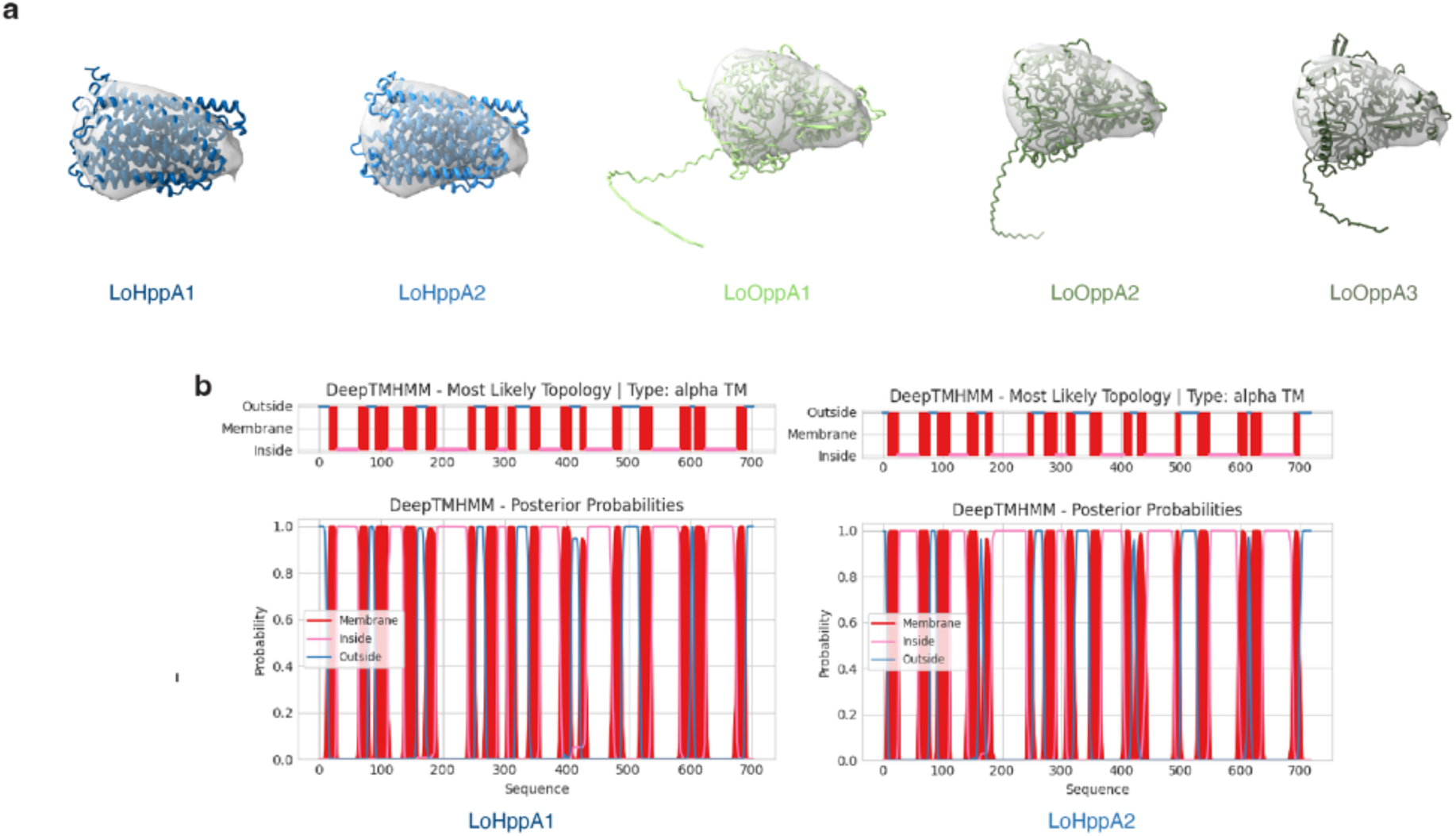
Structural docking of candidate surface proteins with the highest overlap scores. **a,** AlphaFold predicted structures of putative pyrophosphate-energized sodium pumps (LoHppA1/2) and putative oligopeptide-binding proteins (LoOppA1/2/3) fitted into the subtomogram average of the SSDs. **b,** Protein topology prediction of LoHppA1 and LoHppA2 using DeepTMHMM. LoHppA proteins are predicted to be multipass transmembrane proteins and do not have a predicted extracellular domain. See Extended Data Fig. 6 for LoOppA1-3.

**Extended Data Fig. 6.**
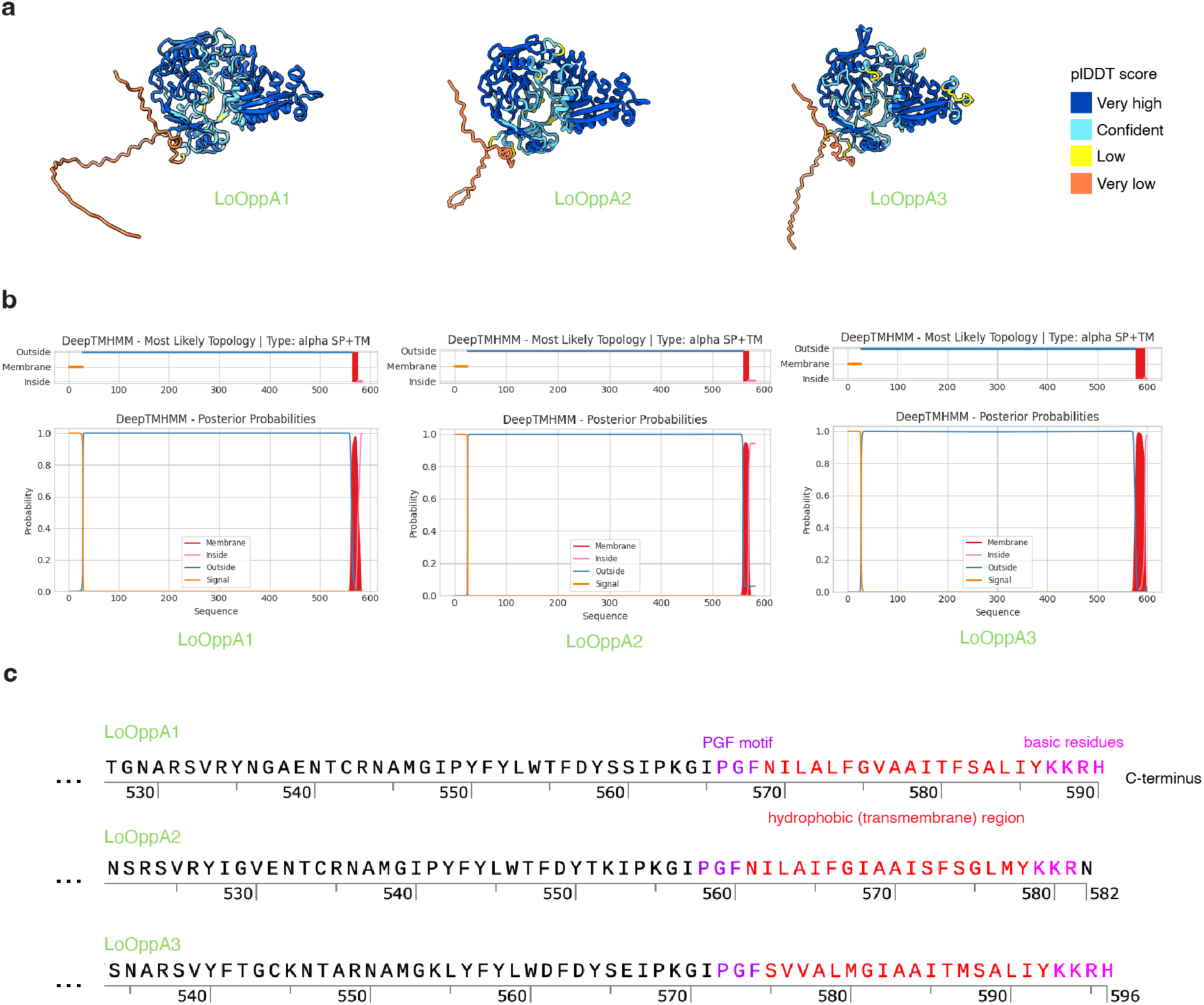
Structure and topology prediction of LoOppA1/2/3 and their sequence features. **a,** AlphaFold predicted structures of the unliganded conformations of LoOppA1/2/3, colored by plDDT score. pTM values of 0.81-0.82 are calculated from each model, indicating high quality and confident predictions. **b,** Protein topology prediction of LoOppA1/2/3 using DeepTMHMM. Each of the proteins is predicted to be a mostly extracellular globular protein that contains an N-terminal signal peptide and a C-terminal alpha helical transmembrane region. **c,** Sequence features of the PGF-CTERM sorting signal^37^ domain found in the C-termini of LoOppA1/2/3.

**Extended Data Fig. 7.**
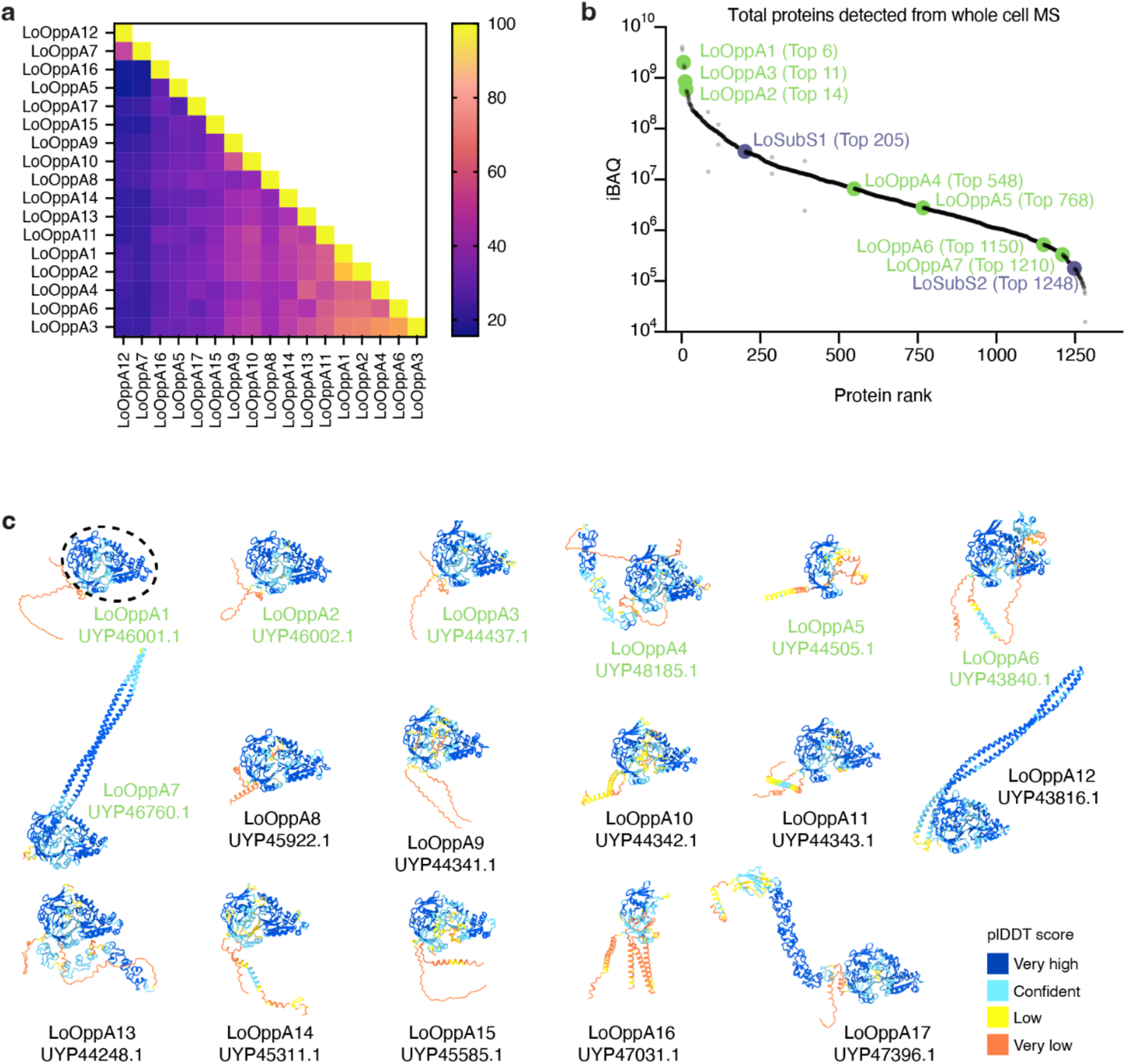
OppA homologs found in the *Ca.* L. ossiferum genome. **a,** Percent identity matrix of LoOppA protein sequences, revealing low sequence identity between different LoOppA homologs. **b,** Protein abundance of different LoOppA homologs shows that only LoOppA1-7 can be detected and with varying abundances. **c,** AlphaFold predicted structures of the open conformations of all LoOppA homologs, colored by plDDT score. Note the conserved oligopeptide-binding region (indicated by dashed line) observed in all predicted structures.

**Extended Data Fig. 8.**
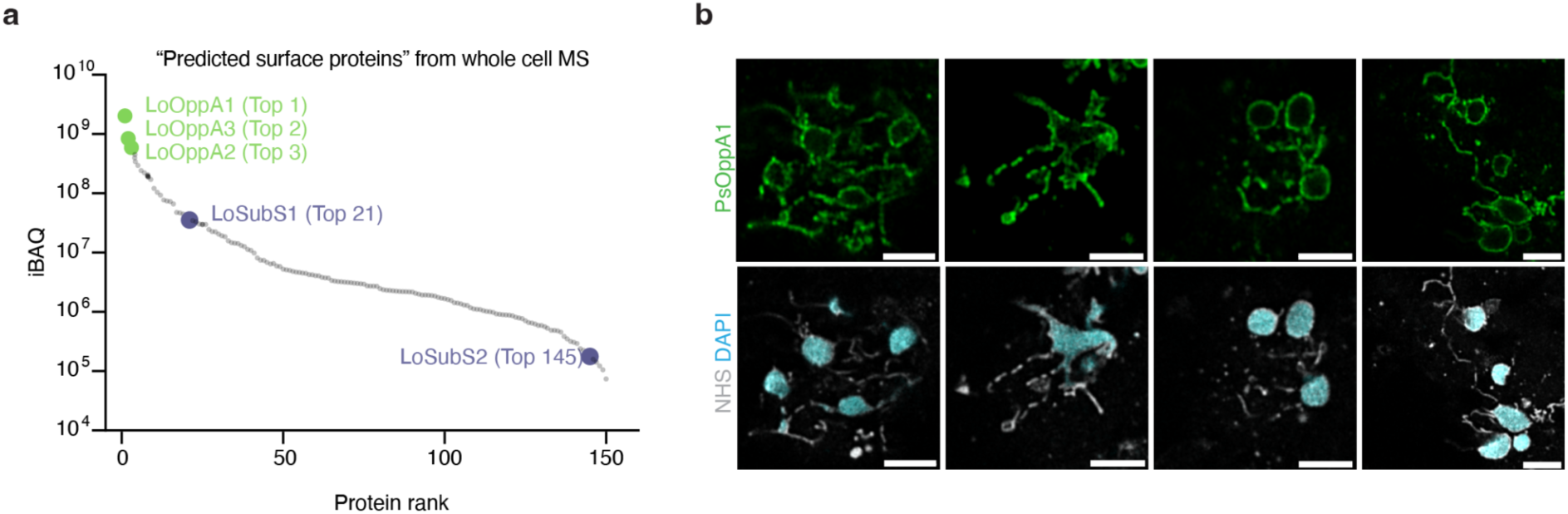
OppA homologs are the major lokiarcheal surface proteins. **a,** Abundances of LoOppA homologs in the whole cell MS-based filtered “predicted surface proteome”, including only proteins predicted to have a signal peptide and/or a transmembrane domain. The dataset identifies LoOppA1/2/3 as the most abundant predicted surface proteins. **b,** Additional examples of U-ExM of *P. syntrophicum* combined with NHS-ester/DAPI staining and post-expansion immunofluorescence staining using PsOppA1 primary antibodies and anti-rabbit IgG secondary nanobodies. Shown are central slices of z-stacks. Scale bars: 1 μm (biological scale).

**Extended Data Fig. 9.**
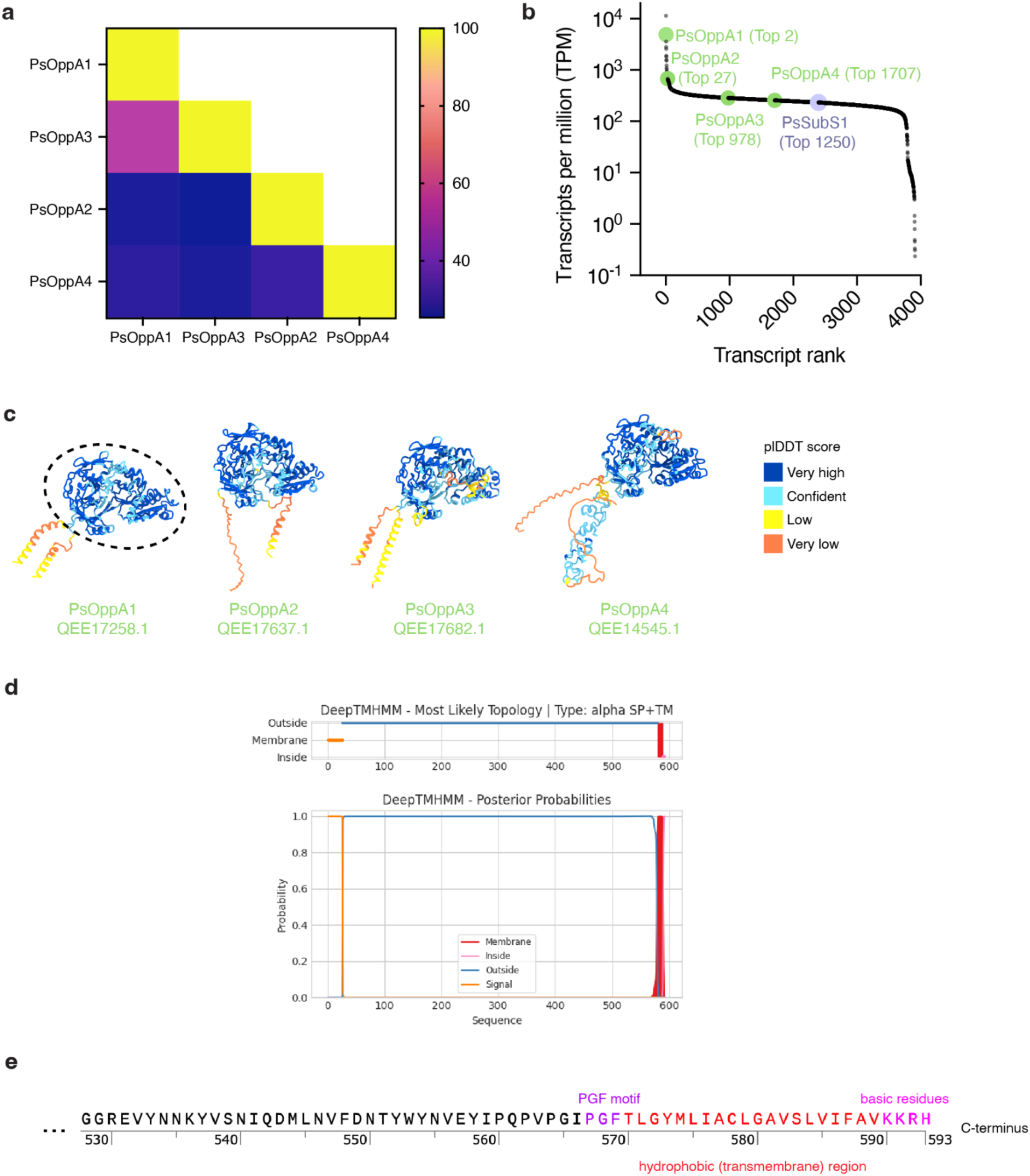
OppA homologs found in the *P. syntrophicum* genome. **a,** Percent identity matrix of PsOppA protein sequences, revealing low sequence identity between different PsOppA homologs. **b,** Transcript abundances of different PsOppA homologs shows transcription of all homologs with varying abundances. **c,** Similar to the LoOppA structures, AlphaFold predicted structures show a conserved oligopeptide-binding region (indicated by dashed line) of PsOppA homologs. **d,** Protein topology prediction of PsOppA1 using DeepTMHMM. PsOppA1 is predicted to be a mostly extracellular globular protein that contains an N-terminal signal peptide with a C-terminal alpha helical transmembrane region. **e,** Sequence features of the PGF-CTERM sorting signal domain found in the C-terminus of PsOppA1.

**Extended Data Fig. 10.**
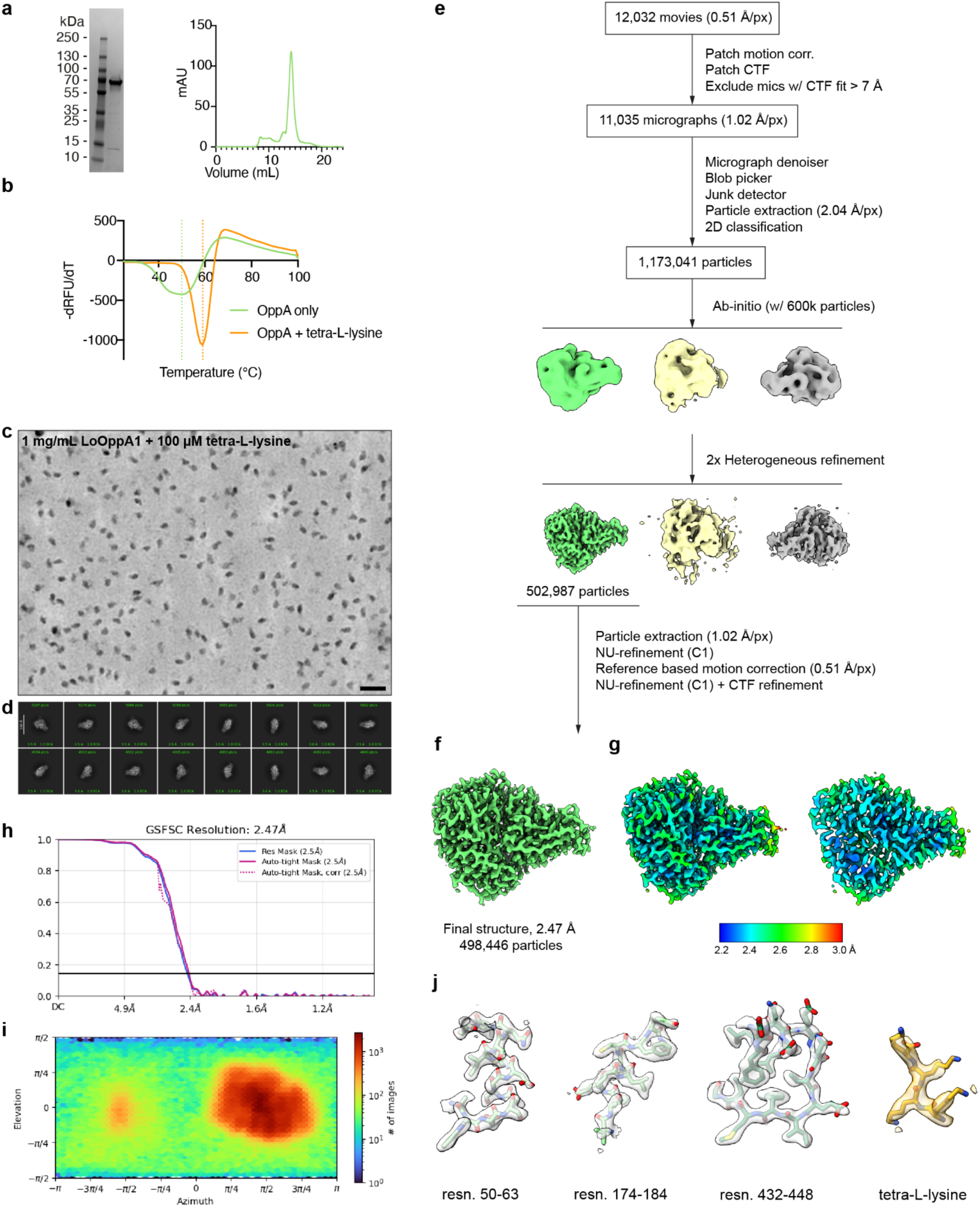
Purification, thermal shift assay of LoOppA1, and cryo-EM data processing workflow of LoOppA1/tetra-L-lysine. **a,** SDS-PAGE gel and size exclusion chromatogram of purified LoOppA1. **b,** Shift in melting temperature of LoOppA1 upon addition of tetra-L-lysine detected using a thermal shift assay. The melting temperature was defined as the temperature corresponding to the maximum rate of change in fluorescence (-dRFU/dT) due to thermal denaturation. **c,** Representative denoised micrograph of LoOppA1/tetra-L-lysine used for cryo-EM data processing. **d,** Representative 2D classes of LoOppA1/tetra-L-lysine. Scale bar: 20 nm. **e,** Cryo-EM data processing workflow for the reconstruction of LoOppA1/tetra-L-lysine. See Materials and methods for details. **f,** Final density map at 2.47 Å, colored in green. **g,** Final density colored by local resolution from 2.2 to 3.0 Å. Both external surface view (left) and a sliced view through the substrate binding pocket (right) is shown. **h,** Gold-standard Fourier shell correlation (GSFSC) for the final reconstruction of LoOppA1/tetra-L-lysine. **i,** Viewing direction distribution plot for the final reconstruction of LoOppA1/tetra-L-lysine. **j,** Representative densities (transparent) of amino acids and their residue numbers (resn). Density of the ligand modeled as tetra-L-lysine is also shown.

**Extended Data Fig. 11.**
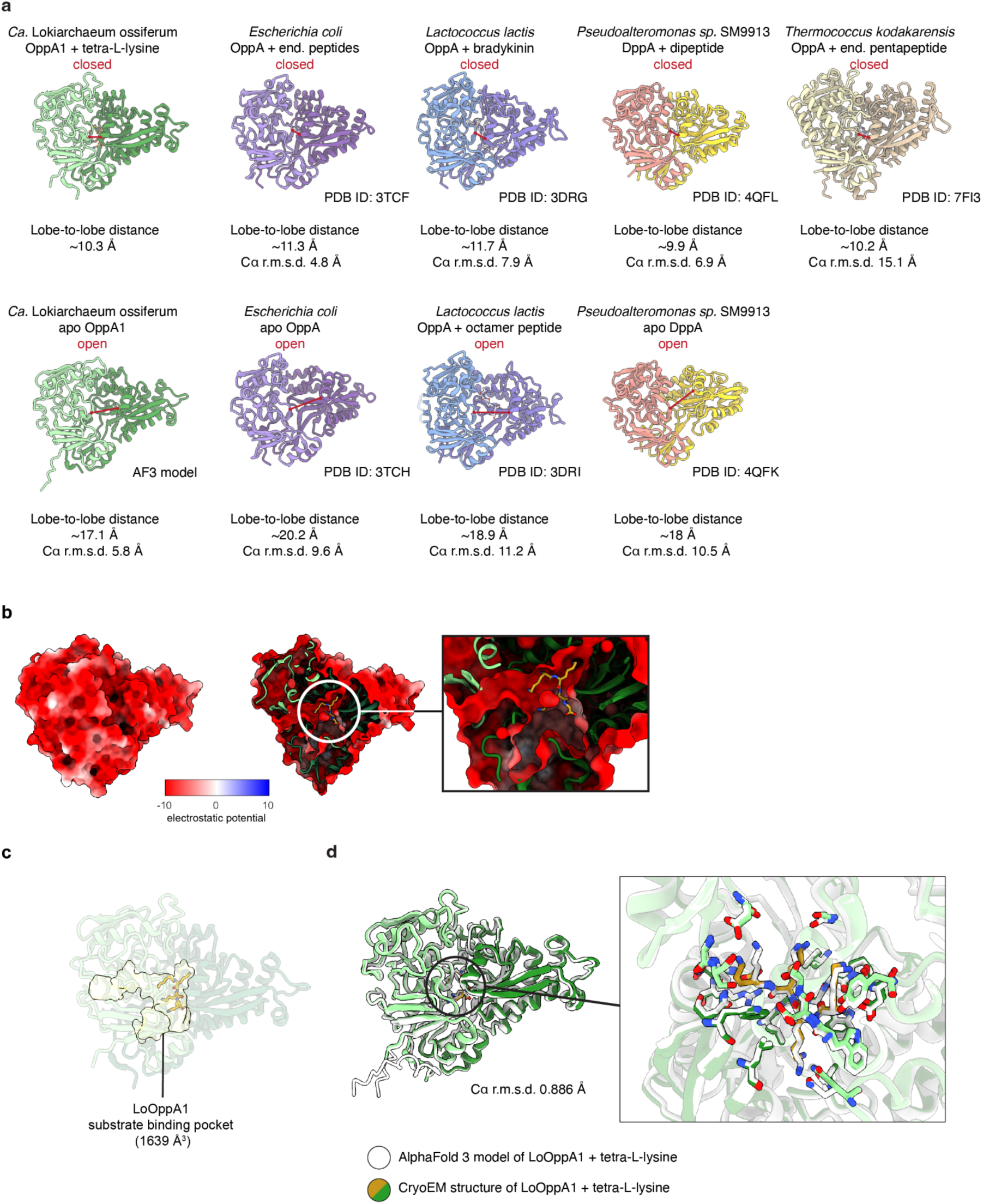
LoOppA1 bound to tetra-L-lysine adopts the closed conformation. **a,** Comparison of the closed and open conformations of LoOppA1 and of homologous proteins from other bacteria and archaea for which structures are available. A model of apo LoOppA1 in its open confirmation was obtained with AlphaFold 3. The structures were compared based on the lobe-to-lobe distance (measured between the Cα atoms of S53 and W430 in LoOppA1 and the equivalent residues in the other structures, shown as a red line) and their Cα root-mean-square deviation from the cryoEM structure of LoOppA1. The short lobe-to-lobe distance in LoOppA1 (∼10.3 Å) indicates that the structure adopts the closed conformation. **b,** Calculated surface electrostatic potential of LoOppA1/tetra-L-lysine. Negative potential is colored red through white to blue for positive potential. External surface view (left), a sliced view (center) and an enlarged view (right) through the substrate binding pocket is shown. Note the highly negative surface electrostatic potential that is favorable for binding of positively charged substrates. **c,** Putative substrate binding cavity size (1,639 Å^3^) of the determined LoOppA1/tetra-L-lysine structure calculated by POCASA^111^. **d,** Comparison of the cryoEM structure (green/orange) and the AlphaFold 3 derived model (white) of LoOppA1 in complex with tetra-L-lysine. A close-up view of the substrate binding pocket is shown on the right. The two structures are nearly identical, as evidenced by the r.m.s.d. value of 0.886 Å.

**Extended Data Fig. 12.**
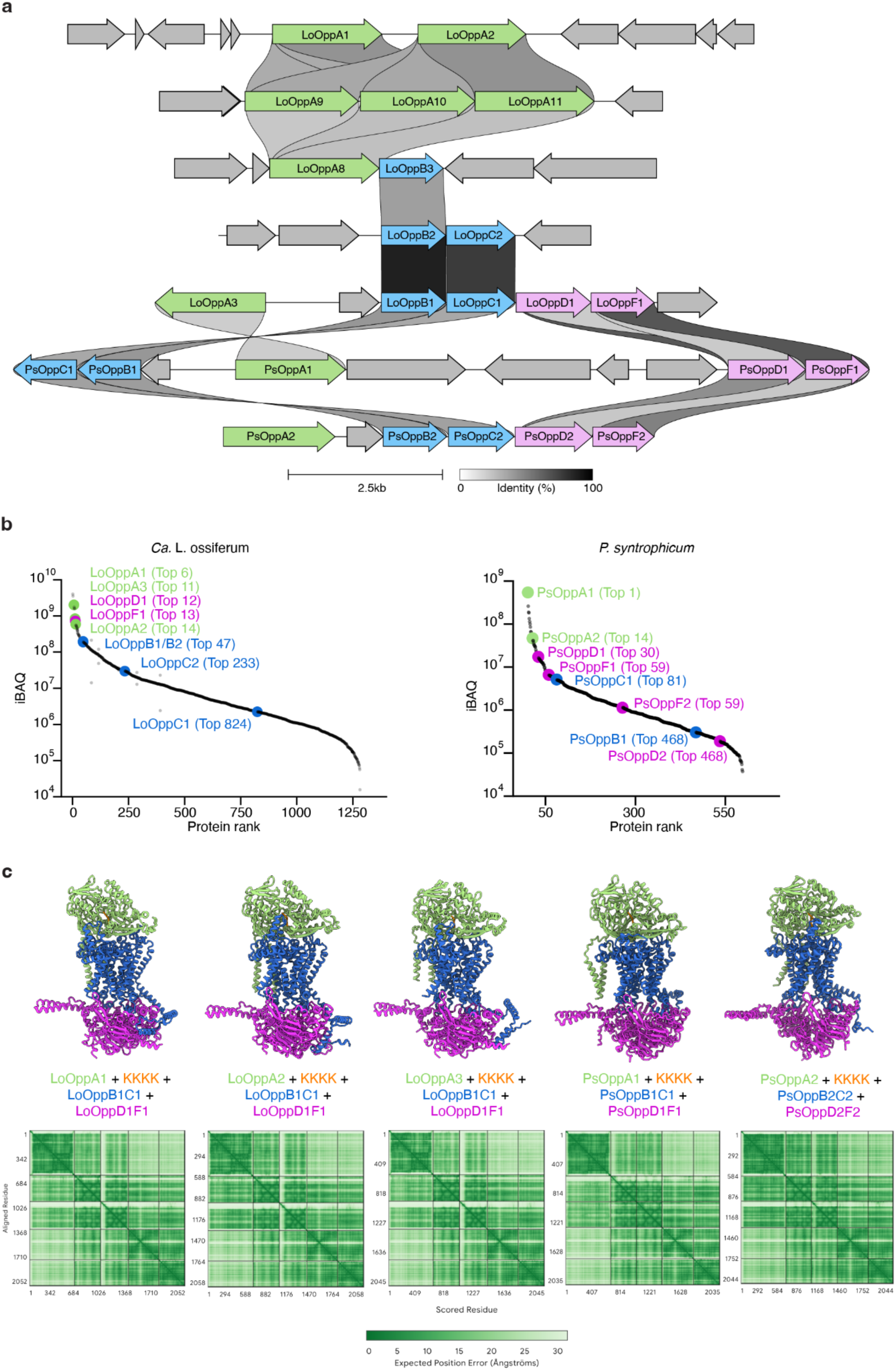
The predicted Opp transporter complexes of *Ca.* L. ossiferum and *P. syntrophicum*. **a,** *opp* operons identified in the genomes of *Ca.* L. ossiferum and *P. syntrophicum*. Only gene clusters containing at least two *opp* genes are shown. The two most highly translated LoOppA homologs Lo*oppA1* and Lo*oppA2* are found colocalized in the genome. The genomic neighborhoods of Lo*oppA3*, Ps*oppA1*, and Ps*oppA2* contain the genes needed for the full complex (*oppABCDF*). **b,** Protein abundances of Opp proteins. The consistency in high abundances of most proteins composing the transporter complex suggests that a high ratio of assembled Opp complexes can be expected. **c,** AlphaFold-multimer predicted structures of the Opp transporter complexes and their predicted aligned error plots (bottom). An ipTM value of 0.72-0.82 was calculated for each multimeric structure, indicating a confident prediction. LoOppA1/2/3 are predicted to form a complex with LoOppB1C1D1F1.

**Extended Data Fig. 13.**
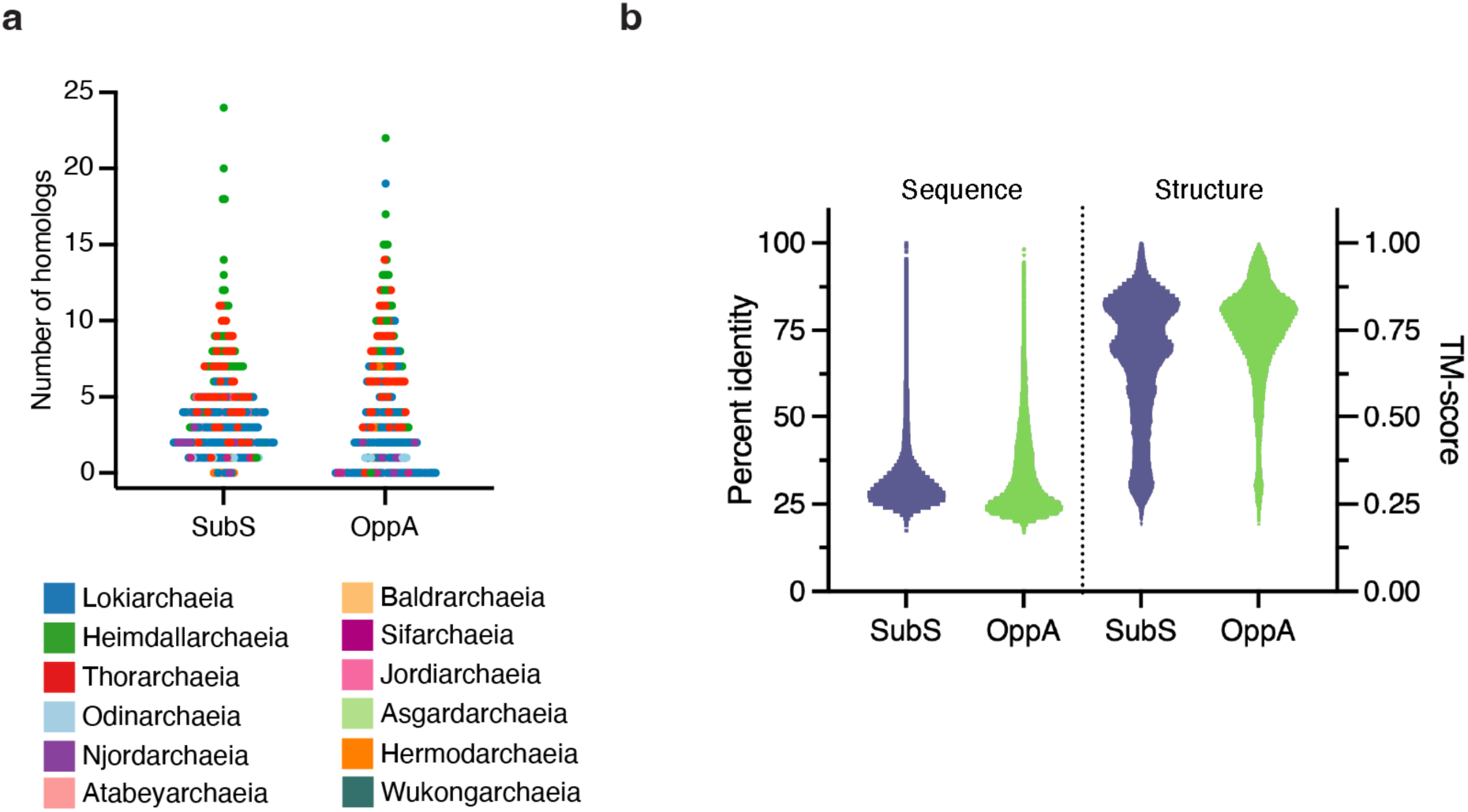
Predicted homologs of surface-anchored SubS and OppA found in other Asgard archaeal genomes. **a,** Number of predicted homologs of SubS and OppA containing a signal peptide and/or transmembrane domains found in other genomes of Asgard archaea. Each dot represents a single genome. **b,** Calculated amino acid sequence identities (left) and predicted structure similarity (right) between identified SubS and OppA homologs. Each dot represents a pairwise comparison between different homologs. The higher the TM-score, the higher the similarity of the structure between two proteins.

**Extended Data Fig. 14.**
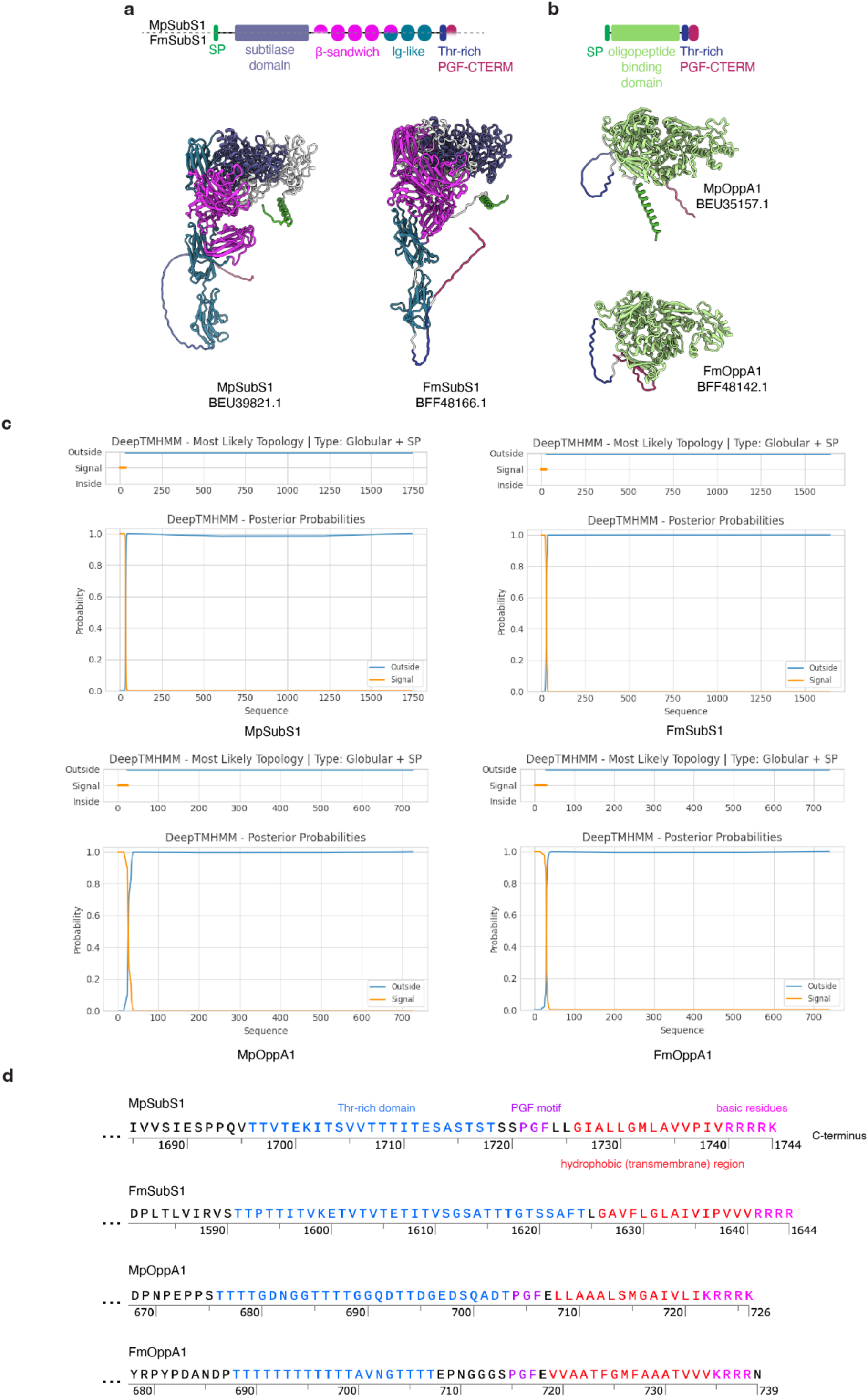
Predicted homologs of surface-anchored subtilase and OppA in cultured hodarchaeal strains *Ca.* M. peptidophila and *Ca.* F. multiprotrusionis. **a,b,** Predicted structures of closest structural homologs identified in Fig. 5 of lokiarchaeal subtilase (**a**) and OppA (**b**) in *Ca.* M. peptidophila and *Ca.* F. multiprotrusionis. Note their similar domain organization to that of the homologs found in the lokiarchaeal strains. **c,** Protein topology prediction of MpSubS1, FmSubS1, MpOppA1, and FmOppA1 using DeepTMHMM. All homologs are predicted to be mostly extracellular globular proteins that contain N-terminal signal peptides. **d,** Sequence features of the PGF-CTERM sorting signal domain found in the C-termini of MpSubS1, FmSubS1, MpOppA1, and FmOppA1.

**Extended Data Fig. 15.**
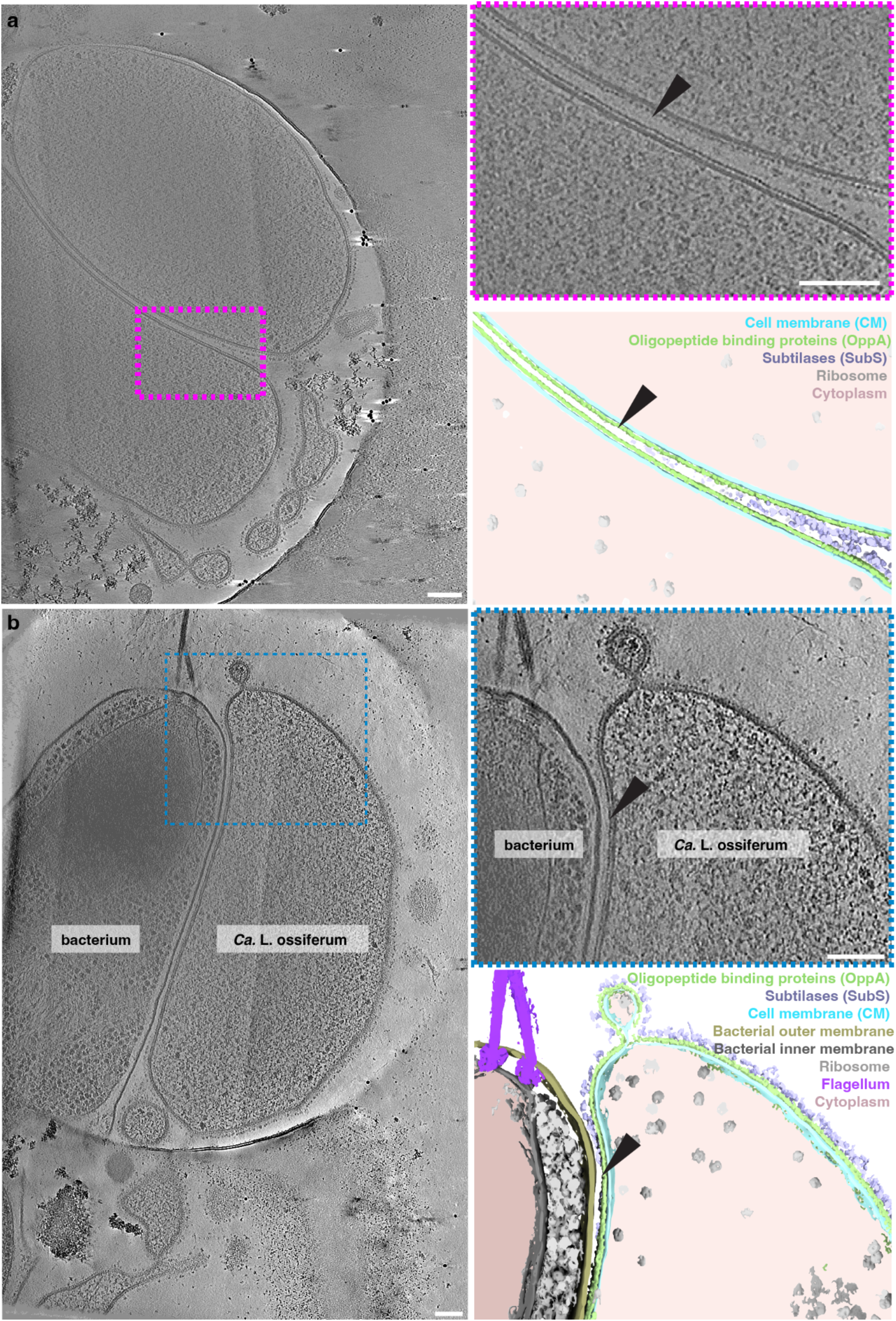
The cell surface is remodeled at cell-cell contact sites. Shown is a slice of a cryo-tomogram between two *Ca.* L. ossiferum cells (a) and *Ca.* L. ossiferum cell with a presumably lysing bacterium (b) with their corresponding segmentations. Note the absence of the LSDs and unchanged distribution of SSDs in regions of close membrane contact indicated by black arrowheads, suggesting remodeling of the cell surface architecture. Scale bars: 100 nm.

