## Supplementary Table 1 for "Visual surfaceomics reveals the substrate-scavenging cell surface of Asgard archaea"

Supplementary Table 1. CryoEM data statistics of LoOppA1 + tetra-L-lysine.

|  | **LoOppA1 + tetra-L-lysine** |
| --- | --- |
| **Data collection and processing** |  |
| Nominal magnification  Voltage (kV)  Electron exposure (e^-^/Å^2^)  Defocus range (µm)  Pixel size (Å/px)  Symmetry Imposed  Final particles (No.)  Map resolution (Å)  FSC threshold  **Refinement**  Map sharpening B factors (Å^2^)  Model composition  Atoms | 165,000  300  ~53  -0.4 to -1.8  0.51 (binning = 1)  C1  498,446  2.47  0.143  -106.3  4190 (Hydrogens: 0) |
| Protein residues  Chains  R.M.S deviations  Bond length (Å)  Bond angles (°)  Validation  MolProbity score  Clashscore  Rotamer outlier (%)  Ramachandran plot  Favored (%)  Allowed (%)  Outlier (%)  Masked CC | 529  2  0.002  0.455  1.07  2.82  0.22  98.67  1.33  0  0.87 |
